# Synthetic tools to redirect the ubiquitin E3 ligase activity of, PRT1, a plant-specific N-recognin

**DOI:** 10.1101/2025.11.27.690864

**Authors:** Keely E. A. Oldham, Suyan Yee, Kai Xun Chan, Peter D. Mabbitt

## Abstract

The Arg/N-degron pathway regulates important agronomic traits. In plants, the PRT1 protein binds to aromatic amino-terminal residues and recruits ubiquitin (Ub) conjugating enzymes to ubiquitinate substrate proteins. Here we demonstrate that PRT1 recruits the UBC35-UEV1A complex via RING1 of PRT1. This stimulates UBC35 to produce K63-linked Ub chains. To afford chemical control over Ub signalling, we construct a synthetic substrate of PRT1 with an N-terminal calmodulin-binding peptide (CBP). The addition of the CBP sequence to the substrate creates an off switch for PRT1-dependent ubiquitination that is regulated by calcium and calmodulin. Finally, we produce a set of nanobodies that are recognised by PRT1 and induce the ubiquitination of a reporter protein. Altogether, we introduce and biochemically validate several new tools to both redirect and chemically control the Arg/N-degron pathway.

## Introduction

The Arg/N-degron pathway regulates important agronomic traits such as submergence tolerance and immunity^1–6^. In plants at least three Ubiquitin E3 ligases (E3s) participate in the Arg/N-degron pathway^2,7^. Two of these, PROTEOLYSIS 6 (PRT6) and BIG are orthologous to mammalian ubiquitin protein ligase E3 component N-recognin 1 (UBR1) and UBR4, respectively^8–10^. Both PRT6 and BIG interact with a single family of ubiquitin conjugating enzymes (UBC1 and UBC2 in *Arabidopsis thaliana*) ^7,9^. The third Arg/N-recognin, known as PROTEOLYSIS 1 (PRT1), appears to be unique to plants ^4,5,10–15^. This makes PRT1 an attractive target for developing plant-specific synthetic biology tools ^2,16,17^. Commercial interest in targeted protein degradation (TPD) tools for both plants and plant pests is increasing, as growers seek to reduce their reliance on non-specific agrichemicals ^16,18,19^.

There is substantial evidence that PRT1 binds to substrates (N-degrons) with aromatic (Phe, Trp, Tyr) amino-termini and that PRT1 recruits a ubiquitin conjugating enzyme (E2), in the UBC8 family, to build Ub chains on those N-degrons^11,13–15,20^. In *A. thaliana* the UBC8 family includes 8 members (UBC8, UBC9, UBC10, UBC11, UBC12, UBC28, UBC29, and UBC30) ^21,22^. Here we investigate whether PRT1 can recruit ubiquitin conjugating enzymes (E2s), other than UBC8 family members, in order to promote the formation of Ub chains with different linkages. Typically, Ub chains are linked by isopeptide bonds between the carboxyl-terminal glycine of Ub and lysine residues (Ub K6, K11, K27, K29, K33, K48, or K63), or by peptide bonds between the carboxyl-terminal glycine of Ub and the amino terminal methionine (Ub M1) ^23^. Ester bonds between the carboxyl-terminal glycine of Ub and the hydroxyl group of serine and threonine residues can also be formed, although the physiological significance of these linkages is unclear ^24^. Here, we find that PRT1 recruits the ubiquitin conjugating enzyme UBC35 to build Ub chains on a model N-degron substrate. We also find that PRT1 interacts with a complex of ubiquitin-conjugating enzyme E2 variant 1A (UEV1A) and UBC35 to build K63-linked Ub chains.

Inspired by naturally occurring N-degrons ^1^, we produce a synthetic reporter substrate with an N-terminal calmodulin-binding peptide (CBP); the addition of the CBP sequence creates a conditional off-switch for PRT1 dependent ubiquitination that is regulated by calcium signalling. Finally, we design a set of nanobodies that are recognised by PRT1 and induce the ubiquitination of a reporter protein. Altogether, we present a set of synthetic biology tools for redirecting the plant Arg/N-degron pathway.

## Results and Discussion

In the absence of a substrate, PRT1 is ubiquitinated by UBC8 (**Fig 1a**) ^11,13,14^. We took a panel of catalytically diverse E2s ^22^ and evaluated their ability to ubiquitinate PRT1 with fluorescent (Cy3-labeled) Ub. We observed ubiquitination with both UBC8 and UBC35 (**Fig 1b**). Given that UBC35 is analogous to yeast UBC13 and that its activity is enhanced by UEV1 proteins (UEV1A, UEV1B, UEV1C, and UEV1D in *A. thaliana*) ^21,25–27^, we assayed PRT1 with UBC35 and UEV1A **(Fig 1c, Supplementary Fig 1**). Our *in vitro* experiments used *A. thaliana* UBC35 and *A. thaliana* UEV1A, but it is important to note that UBC35 and UBC36 are highly similar to one another and are functionally redundant in *A. thaliana*^26,27^. In a PRT1 autoubiquitination assay both UBC35-UEV1A and UBC35-UEV1C enhanced the level of PRT1 ubiquitination relative to UBC35 alone (**Supplementary Fig 1)**. PRT1 enhanced the rate of di-ubiquitin (Ub_2_) chain formation by UBC35-UEV1A (**Fig 1c, d**). We observed a similar rate enhancement with UBC35-UEV1C (**Supplementary Fig 2**). The unanchored Ub dimers formed by UBC35 are likely to be K63 linked, as no chains were formed when UBC35 was assayed with K63R Ub (**Fig 1e, f**).

**Fig 1.**
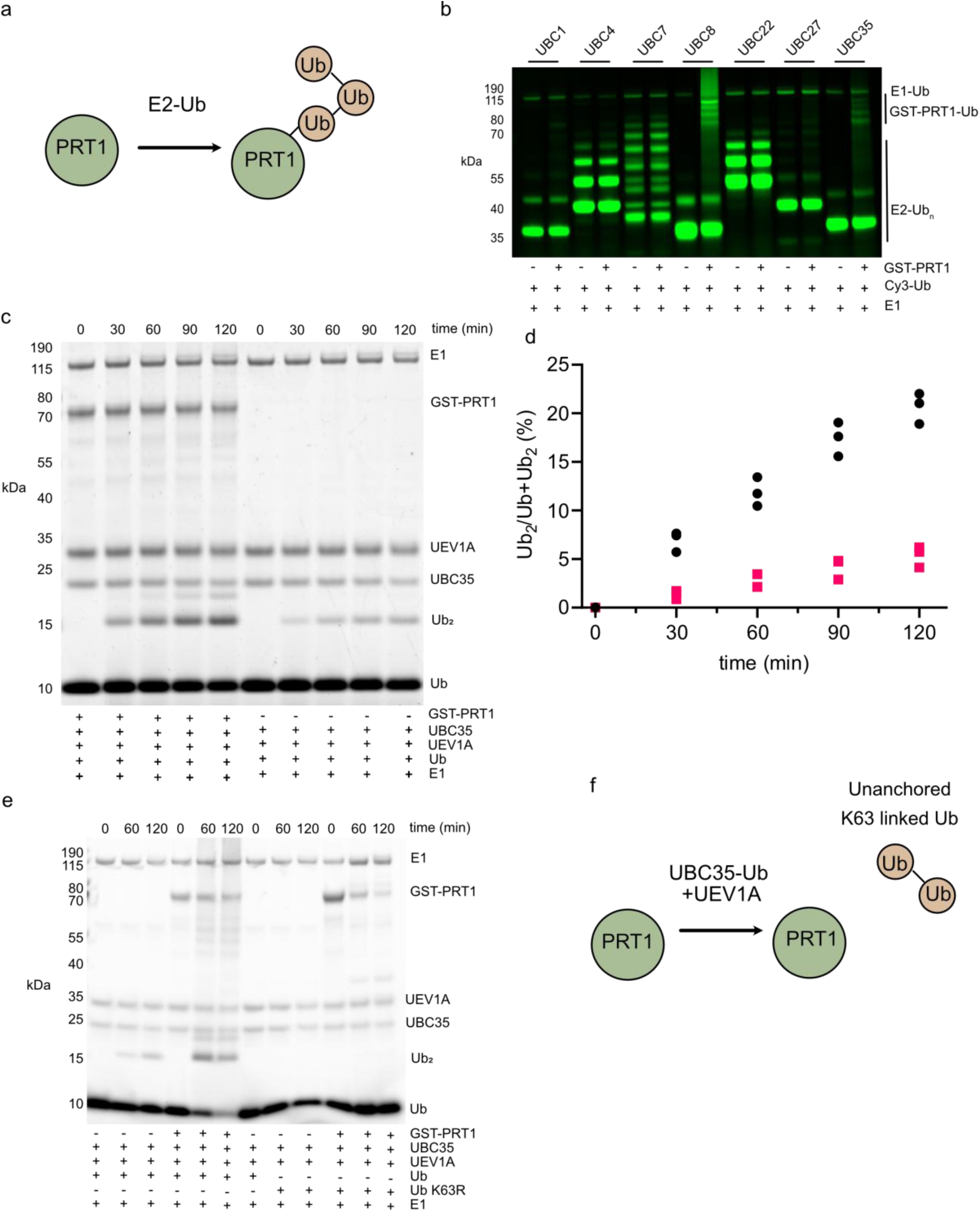
PRT1 interacts with both UBC8 and UBC35. **a** Schematic of PRT1 ubiquitination by an E2. **b** The indicated E2s were incubated with GST-PRT1, ATP, E1, and fluorescently labelled (cy3) Ub for 3 h. **c** GST-PRT1 was incubated with ATP, E1, UBC35, UEV1A and Ub for the indicated times (representative Coomassie blue stained SDS-PAGE gel). **d** Quantification of di-ubiquitin (Ub_2_) in panel c (n=3); circles (with PRT1) and squares (without PRT1). **e** The UBC35-UEV1A complex produces di-ubiquitin (Ub_2_) from Ub but not Ub K63R. GST-PRT1 was incubated with ATP, E1, UBC35, UEV1A and Ub or Ub K63R for the indicated times. **f** Schematic of PRT1 promoting UBC35-UEV1A catalysed formation of K63 linked Ub dimers.

We next assayed the PRT1-E2 complexes with a model N-degron substrate. We modified a previously reported substrate^13^, *Escherichia coli* flavodoxin, so that following Tobacco Etch Virus (TEV) protease-cleavage the protein had a Tyr amino-terminus and a single surface exposed lysine residue. We named this substrate Flav1K (**Fig 2a**). The Flav1K protein has three lysine to arginine substitutions (Lys32Arg, Lys76Arg, and Lys105Arg), relative to the reference sequence (Protein Data Bank (PDB) accession 2M6R). Typical substrates have multiple surface exposed lysine residues, and it is non-trivial to distinguish between multi-mono ubiquitination and Ub chain formation. As Flav1K has a single lysine residue, all isopeptide linked poly-Ub adducts on Flav1K are Ub chains. Henceforth, we refer to the addition of the first Ub to a substrate as priming. We refer to the addition of subsequent Ub moieties as chain extension. As has been previously demonstrated^11,13^, PRT1 directs UBC8 to ubiquitinate substrates with aromatic N-termini (**Fig 2b**). We observed UBC8 dependent ubiquitination of Flav1K with WT, K48R, and K63R Ub. We consistently saw less ubiquitin chain formation when PRT1 was assayed with K63R Ub. These results suggest that PRT1 can promote the formation of multiple Ub chain topologies. We next investigated whether UBC35-UEV1A could produce Ub chains on Flav1K. When PRT1 and Flav1K were incubated with UBC35-UEV1A we did not observe Ub chains on Flav1K (**Fig 2c**). This may be due to UBC35-UEV1A preferentially catalysing the formation of Ub chains that are not linked to FLAV1K. Consistent with this, UBC35 (without UEV1A) transferred Ub to Flav1K (**Fig 2d**). The amount of Ub transferred by UBC35 to FLAV1K was lower than that transferred by UBC8.

**Fig 2.**
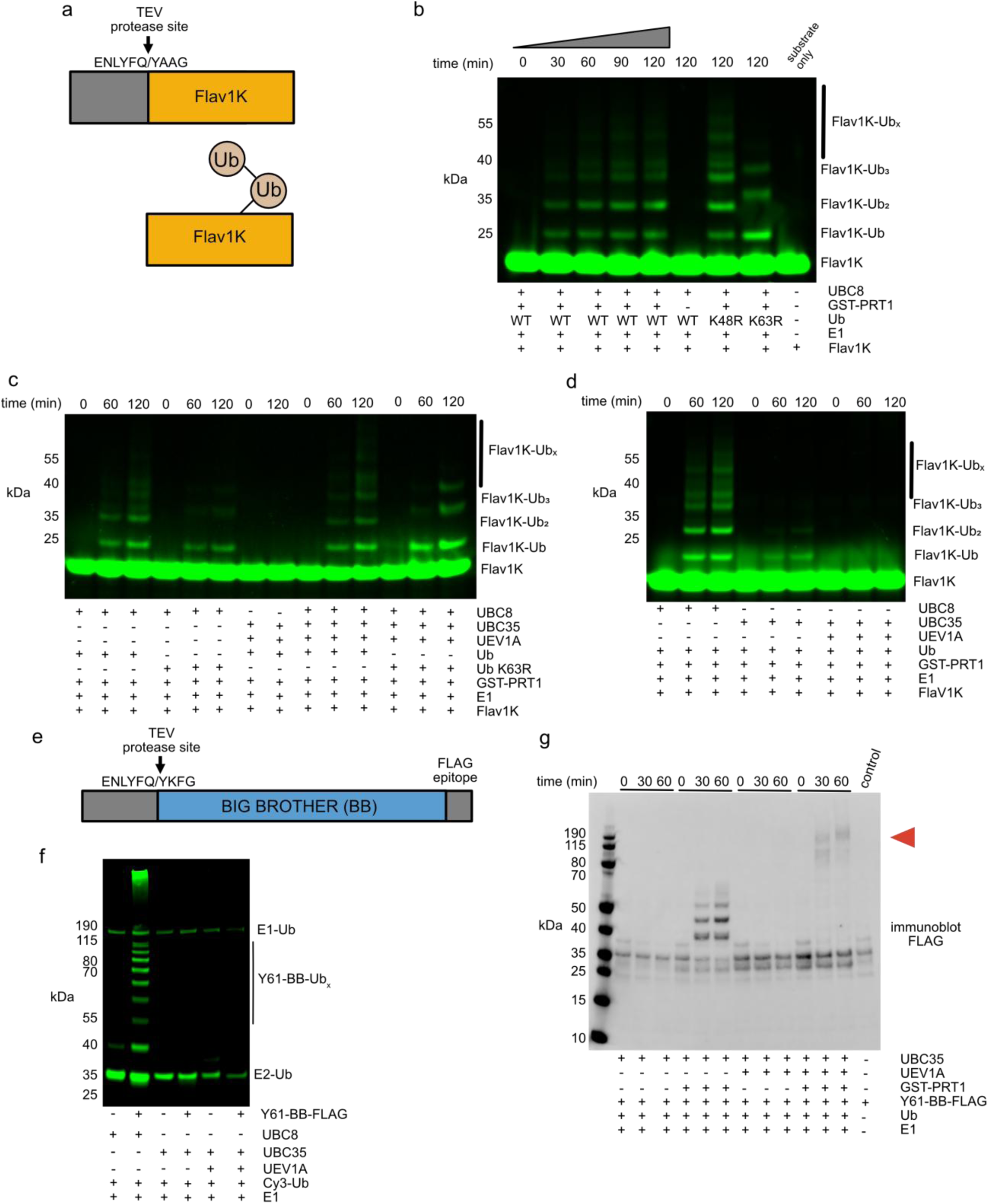
PRT1 directs UBC8 to produce Ub chains on N-degron substrates. **a** Schematic of a model N-degron substrate, following cleavage with TEV protease the amino-terminal Tyr (Y) of engineered flavodoxin (Flav1K) is recognised by PRT1. The Flav1K protein has a single Lys (K) to which ubiquitin may be attached. The single cysteine of FlaV1K was labelled with Cy3. **b** PRT1 directs UBC8 to transfer Ub to Flav1K. GST-PRT1, ATP, E1, UBC8 and fluorescently labelled Flav1K were incubated with the indicated Ub mutants. **c** PRT1 directs UBC8, but not the UBC35-UEV1A complex, to transfer Ub to Flav1K. GST-PRT1, ATP, E1, UBC8, UEV1A, UBC35 and fluorescently labelled Flav1K were incubated with the indicated Ub mutants. **d** PRT1 directs UBC35, in the absence of UEV1A, to transfer Ub to Flav1K. GST-PRT1, ATP, E1, UBC8, UEV1A, UBC35 and fluorescently labelled Flav1K were incubated with wild-type Ub. **e** Schematic of the E3 ligase BIG BROTHER (BB), in *A. thaliana* cleavage of BB results in a Tyr N-terminus (Y61) that is recognised by PRT1. In our recombinant system, BB has an N-terminal TEV protease site and a C-terminal FLAG epitope. Cleavage of the recombinant protein results in Y61-BB-FLAG. **f** In an autoubiquitination assay Y61-BB-FLAG was incubated with ATP, E1, Cy3-Ub, and the indicate E2s for 1 hr. **g** PRT1 directs Ub transfer from UBC35 to Y61-BB. The Y61-BB protein was incubated with ATP, E1, Ub, UBC35, UEV1A and GST-PRT1 for the indicated time. Samples were resolved by SDS-PAGE and immunoblotted with Anti-FLAG antibody (n=3). Red arrow indicates putative high-molecular weight Y61-BB ubiquitin adducts.

The best characterised substrate of PRT1 is an E3 ligase known as BIG BROTHER (BB)^20^. Proteolytically processed BIG BROTHER has an N-terminal Tyr (Y61-BB). The BB protein substrate has 10 lysine residues so a reaction with 3 µM of denatured BB would present 30 µM of substrate-lysine^11^. We produced recombinant Y61-BB with an authentic amino-terminus and a carboxyl-terminal FLAG epitope (**Fig 2e**). There is evidence that BB is a RING E3 ligase and that Y61-BB retains activity with the UBC8 family E2s *in vitro*^11,20,28^. Under our assay conditions Y61-BB activated UBC8 (**Fig 2f**). However, we did not observe Y61-BB autoubiquitination when assayed with UBC35 (**Fig 2f, g**). When Y61-BB was incubated with PRT1 and UBC35 we observed ubiquitination of Y61-BB (**Fig 2g**). Similarly, when Y61-BB was incubated with UBC35-UEV1A and PRT1 we observed high-molecular weight Y61-BB adducts (**Fig 2g**). This data is consistent with PRT1 recruiting the UBC35-UEV1A complex to form poly-ubiquitin chains on Y61-BB.

To better understand the PRT1-substrate interaction we assayed the complex with a larger model substrate, CBP-RFP. The CBP-RFP protein is a lysine-rich substrate with an aromatic (Tyr) N-terminus followed by a calmodulin binding peptide (CBP) and mCherry (RFP) (**Fig 3a**) ^29,30^. In experiments with Flav1K the substrate concentration and substrate-lysine concentration were ∼3.4 µM, whereas in experiments with CBP-RFP the substrate concentration and substrate-lysine concentration were ∼5 µM and 145 µM, respectively. For the N-degron pathway this distinction is important because each degron has a single binding site (N-terminus) but a different number of E2 accessible lysine residues. We observed that PRT1 directed both UBC8 and UBC35 to ubiquitinate CBP-RFP (**Fig 3b**). Altogether our data suggests that PRT1 both stimulates the UBC35-UEV1A complex to produce unanchored K63-linked Ub chains and directs UBC35 to attach Ub to N-degron substrates, including Y61-BB. These data also show that the PRT1-UBC8 complex can ubiquitinate multiple substrates. In further *in vitro* assays with CBP-RFP and other engineered proteins we used UBC8 as it gave a clearer signal than UBC35 for quantification.

**Fig 3.**
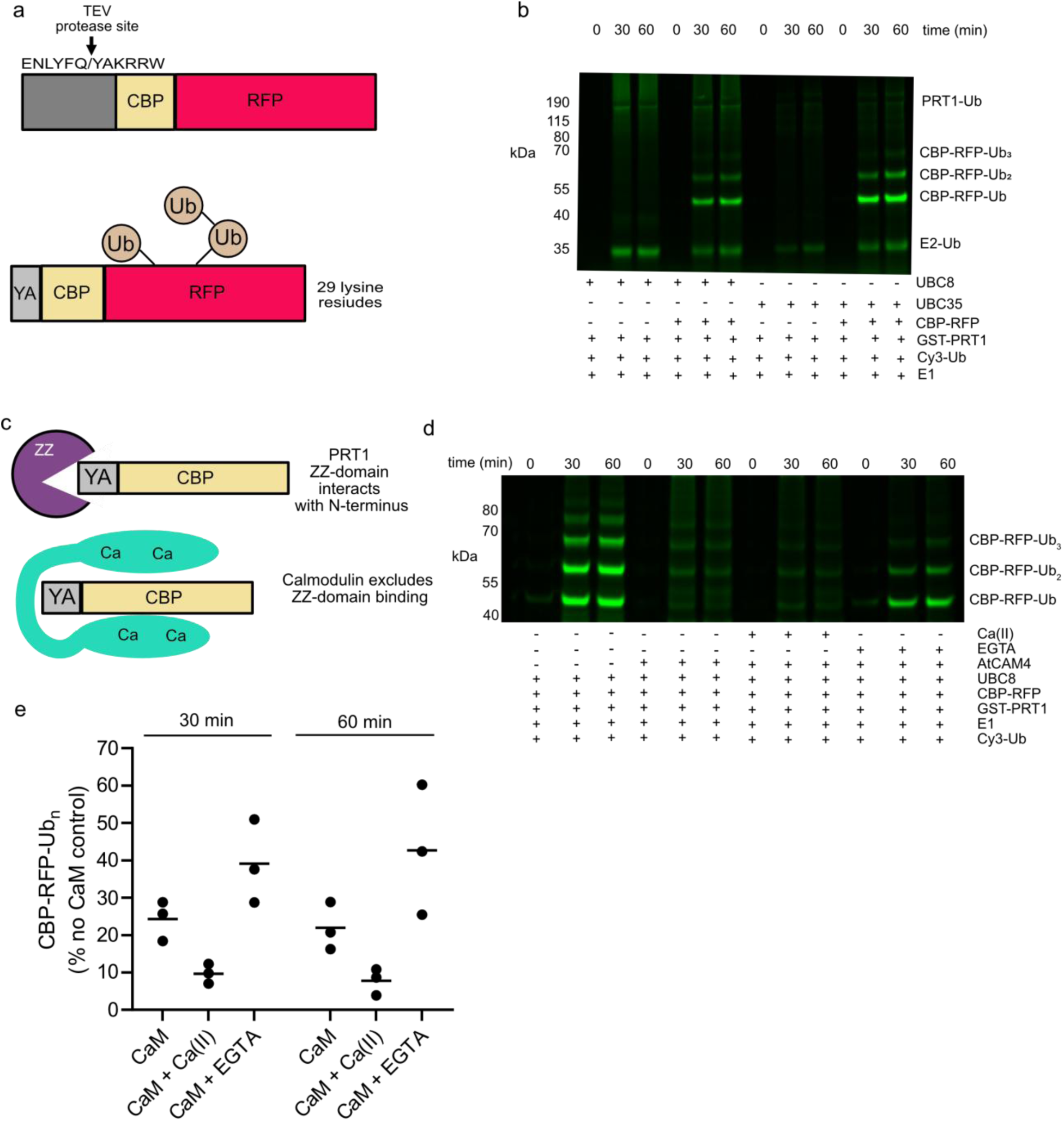
Calmodulin regulated substrate ubiquitination. **a** Schematic of CBP-RFP. A fusion protein with a TEV protease site followed by a calmodulin binding peptide (CBP) and mCherry (RFP). Cleavage of the CBP-RFP results in a tyrosine (Y) N-terminus. The CBP-RFP has multiple surface exposed lysine residues to which Ub may be attached. **b** PRT1 directs UBC8 and UBC35 to transfer Ub to CBP-RFP. GST-PRT1, ATP, E1, UBC8, UBC35 and Cy3-Ub were incubated for the indicated time. The putative CBP-RFP-Ub bands have been labelled. The greater than 80 KDa ubiquitin adducts may be a mixture of CBP-RFP-Ub_x_ and GST-PRT1-Ub_x_. **c** Calmodulin (CaM) and the ZZ-domain of PRT1 compete for an N-terminal calmodulin-binding peptide (CBP). The amino terminus of the CBP-RFP construct has Tyr-Ala (YA) immediately preceding CBP. **d** PRT1 directs Ub transfer from UBC8 to CBP-RFP (5µM), the addition of excess CaM (20 µM) or CaM and 5 mM CaCl_2_ attenuates Ub transfer. Addition of EGTA (5 mM) partially restores Ub transfer (representative gel, n=3). GST-PRT1, CBP-RFP, ATP, E1, UBC8, Cy3-Ub, and AtCAM4 were incubated for the indicated time. **e** Quantification of CBP-RFP-Ub in panel d, the mean is indicated with a bar.

Given that the substrate binding, ZZ-domain, of PRT1 interacts specifically with N-termini^11^, we reasoned that calmodulin (CaM) binding to the CBP region of CBP-RFP would block PRT1 dependent ubiquitination of CBP-RFP (**Fig 3 c**). In the CBP-RFP construct the N-terminus is Tyr, followed by Ala and the CBP sequence (**Fig 3 c**). Experimental evidence suggests that the amino-terminus of the protein will be buried within the ZZ-domain of PRT1^11^. This would sterically exclude other proteins from binding the amino-terminus. Likewise, calmodulin binding to CBP would sterically exclude the amino-terminal region of CBP^30^. In isolation calmodulin has low nanomolar affinity (∼2 nM *K_D_*) for the CBP sequence, this is at least a thousand-fold greater than the affinity of the isolated ZZ-domain for N-termini^11,30^. We observed that the addition of calmodulin (*A. thaliana* CAM4; AtCaM4) attenuated PRT1 dependent ubiquitination of CBP-RFP. As would be expected, the addition of calcium and calmodulin has an additive effect. The action of CaM could be partially reversed by the addition of the calcium chelator EGTA (**Fig 3 d, e**).

We next sought to validate the CBP-RFP reporter *in vivo*. We transiently expressed CBP-RFP, with a stabilising (Met) or destabilising (Tyr) N-terminus, in *A. thaliana* protoplasts and *Nicotiana benthamiana* leaves. Fusion proteins were expressed with an N-terminal Ub, where cleavage of Ub is expected to result in CBP-RFP with either Met or Tyr preceding the CBP sequence^31^ (**Fig 4a**). Under steady state conditions in transformed *A. thaliana* protoplasts, we observed higher RFP fluorescence associated with the stabilising (Met) N-terminus than the destabilising (Tyr) N-terminus. High-resolution confocal microscopy further revealed the subcellular compartment where the majority of protein turnover occurred (**Fig 4b**). Fluorescence from Met-CBP-RFP was observed in both the nucleus and cytosol. Conversely, fluorescence from Tyr-CBP-RFP was relatively low in the cytosol, with some nuclear RFP fluorescence remaining. The nuclear RFP fluorescence functioned as a transformation control. The tendency of untargeted GFP and RFP based reporters to accumulate in the nucleus has been observed by others^32^. For quantification of RFP, we only analysed protoplasts with an RFP signal in the nucleus (**Fig 4b**). Quantification of RFP fluorescence indicated higher RFP fluorescence associated with the stabilising (Met) N-terminus than the destabilising (Tyr) N-terminus (**Fig 4c**). This trend was also observed in transiently transformed *N. benthamiana* leaves (**Fig 4d, e**). Notably, Met-CBP-RFP expression was observed in both the nucleus and cytosol, whereas Tyr-CBP-RFP expression was largely confined to the nucleus (**Fig 4d**).

**Fig. 4.**
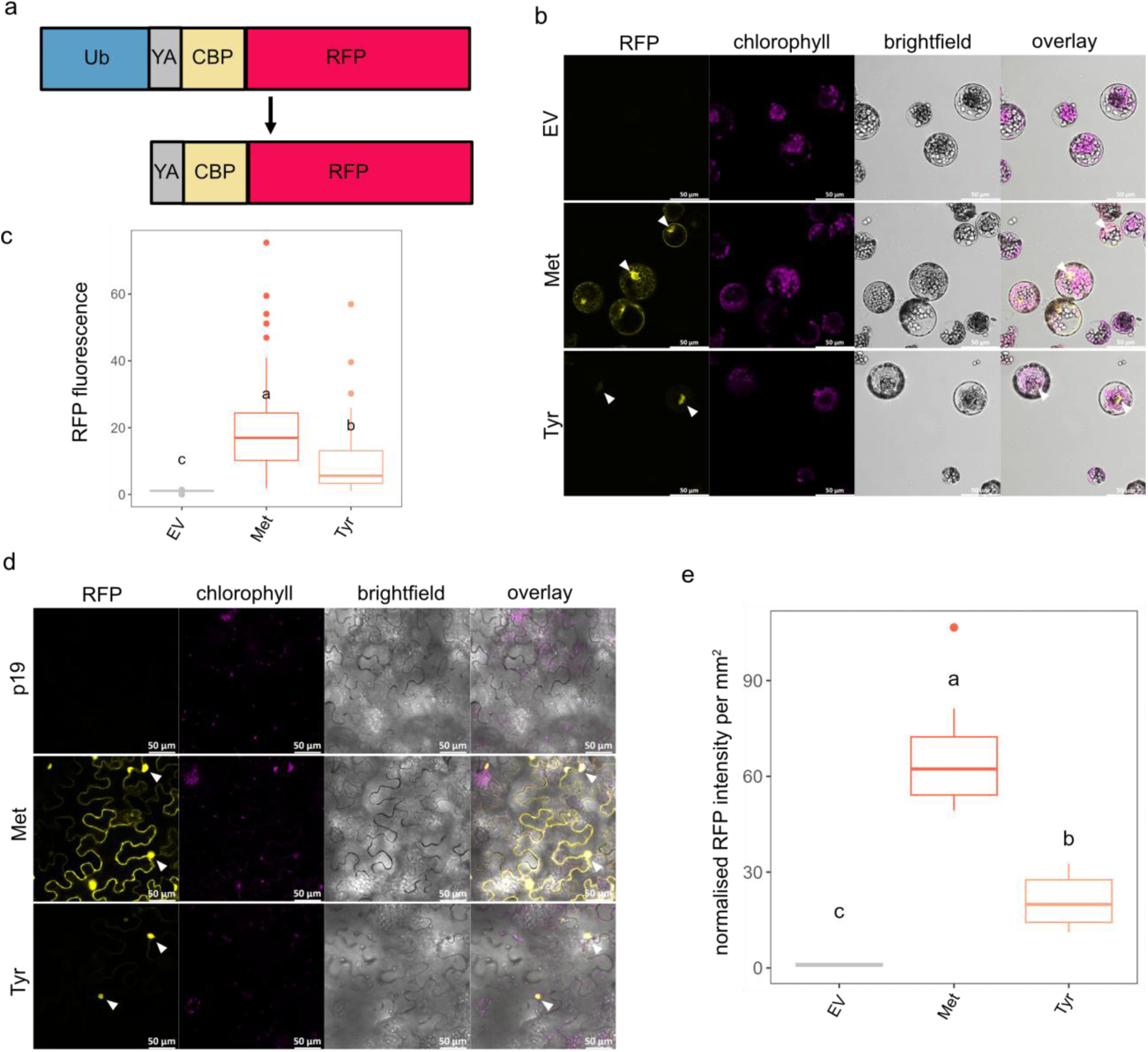
Transient expression of CBP-RFP in *Arabidopsis thaliana* protoplasts and *Nicotiana benthamiana* leaves. **a** Schematic of the fusion proteins transiently expressed *in vivo*. The ubiquitin (Ub) is proteolytically removed from the N-terminus resulting in a Tyr-Ala (YA) of Met-Ala (MA) N-terminus. The calmodulin binding peptide (CBP) and mCherry (RFP) regions are indicated. **b** Representative confocal images of *A. thaliana* protoplasts transiently expressing Met-CBP-RFP, Tyr-CBP-RFP, or an empty vector (EV) control. Images were collected at one day post-transfection (dpt). RFP fluorescence is false-coloured yellow, while chlorophyll autofluorescence is denoted in magenta. White arrows highlight the nuclei within transformed protoplasts. Scale bars = 50 µm. **c** RFP fluorescence quantified in *A. thaliana* protoplasts and plotted as box and whiskers (EV n=70, Met-CBP-RFP n=117, Tyr-CBP-RFP n=59). Lowercase letters (a, b, c) indicate statistically significant differences between reporter constructs, as determined via one-way ANOVA with Tukey’s HSD test. For all comparisons the adjusted *p* values are less than 0.001. **d** Representative confocal images of Met-CBP-RFP and Tyr-CBP-RFP proteins transiently expressed in *N. benthamiana* leaves compared to the p19 negative control. Images were collected at three days post-infiltration (dpi) and are coloured as they are in panel b. White arrows highlight nuclei within transformed epidermal pavement cells. **e** RFP fluorescence normalised to infiltrated leaf area was quantified in 8 transiently transformed *N. benthamiana* leaves. Measurements were plotted as box and whiskers. Lowercase letters (a, b, c) denote statistically significant differences from a one-way ANOVA with Tukey’s HSD test (comparisons and adjusted *p* values: Tyr-p19 *p*=0.008, all other comparisons *p*< 0.001).

To down-regulate components of the N-degron pathway we designed artificial microRNAs (amiRNAs)^33^ cloned as pre-amiRNAs that were embedded, as introns, within the CBP-RFP reporters (**Fig 5a**, **Supplementary Fig 3)**. In this system, presence of RFP fluorescence indicates a transformed cell or protoplast with correctly spliced out pre-amiRNAs. This design was initially assessed in *N. benthamiana* using the monomeric yellow fluorescence protein (mCitrine) as a reporter. An ami-mCitrine sequence was cloned within the intron of CBP-RFP and this construct was transiently co-infiltrated with a construct encoding mCitrine. Expression of RFP with concurrent silencing of the co-expressed mCitrine construct indicated correct splicing of pre-ami-mCitrine from RFP and functional silencing of mCitrine by mature ami-mCitrine **(Supplementary Fig 3).** Consequently, we established ami-mCitrine as an appropriate amiRNA control for the subsequent silencing assays. While the absolute levels of RFP fluorescence were lower when the ami-mCitrine was present, the higher level of Met-CBP-RFP compared to Tyr-CBP-RFP was maintained (**Fig 5b,c, d, Supplementary Fig 4**). The subcellular localisation patterns of RFP fluorescence in the ami-mCitrine controls (**Supplementary Fig 3, Supplementary Fig 4**) also reflect those previously observed in the earlier constructs (**Fig 4b**), where Tyr-CBP-RFP is degraded in the cytosol but still detectable in the nucleus.

**Fig. 5.**
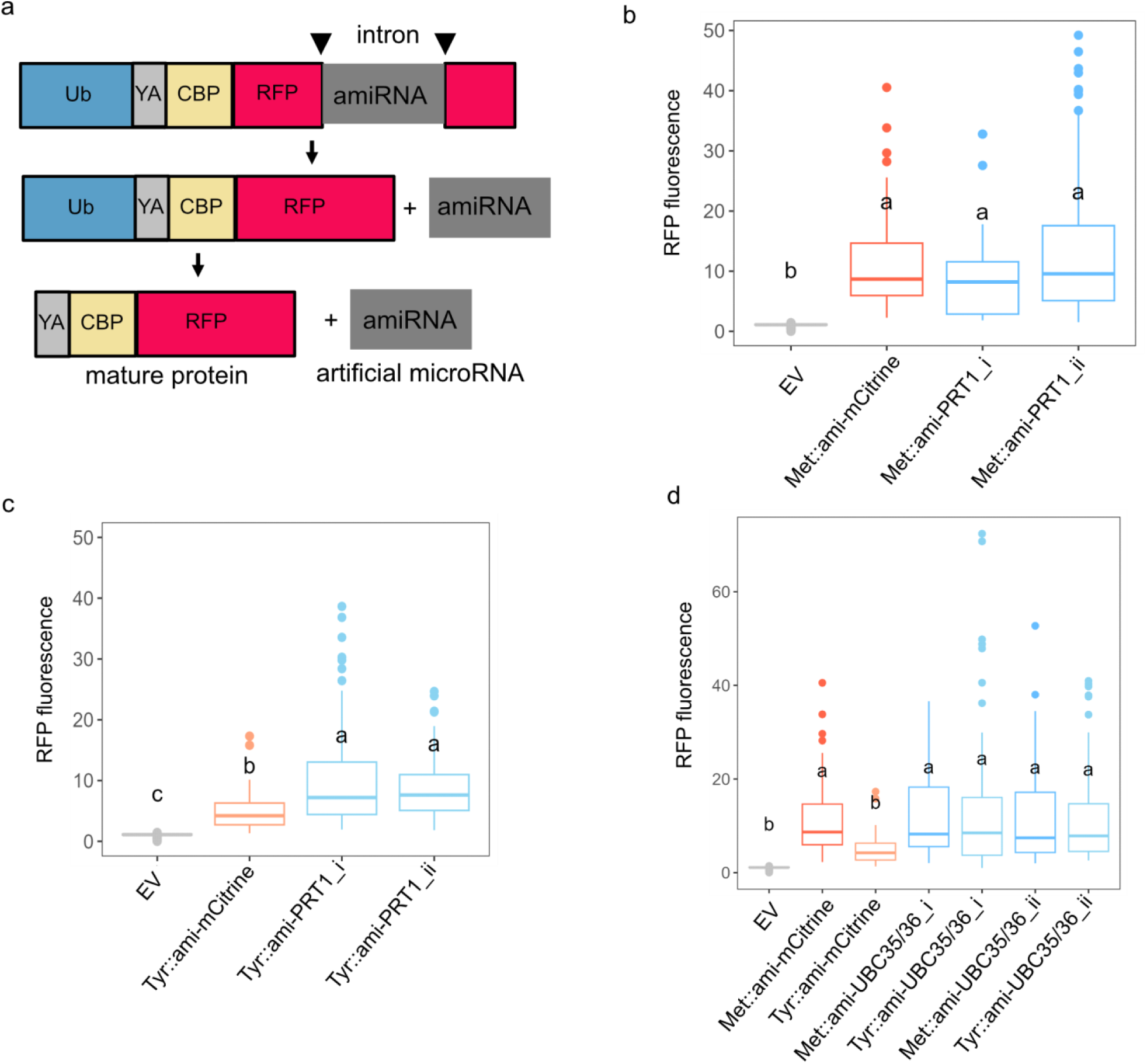
Silencing PRT1 increases the stability of an aromatic N-degron in *Arabidopsis thaliana* protoplasts. **a** Schematic of Ub-CBP-RFP fusion with an artificial microRNA (amiRNA) present as an intron. Correct splicing of the exons is required for the translation of the fluorescent reporter protein. Abbreviations; Ubiquitin (Ub), Tyr-Ala (YA), calmodulin binding peptide (CBP), and RFP (mCherry). **b** Two amiRNAs targeted to PRT1 (ami-PRT1_i or ami-PRT1_ii), or a control amiRNA targeted to mCitrine (ami-mCitrine), were encoded within the CBP-RFP reporter. The reporter protein had a stabilising (Met) N-terminus. Confocal images were collected one day post-transfection and RFP fluorescence was quantified and plotted as box and whiskers (Empty vector (EV) n= 70, Met::ami-mCitrine n=61, Met::ami-PRT1_i n=36, Met::ami-PRT1_ii n=114). Lowercase letters (a, b) denote statistically significant differences from a one-way ANOVA with Tukey’s HSD test. Constructs sharing the same letter are not significantly different. For all significant comparisons, the adjusted *p* values are less than 0.001. **c** As panel b, except the reporter has a destabilising (Tyr) N-terminus (EV n=70, Tyr::ami-mCitrine n=42, Tyr::ami-PRT1_i n=94, Tyr::ami-PRT1_ii n=90). Lowercase letters (a, b,c) denote statistically significant differences from a one-way ANOVA with Tukey’s HSD test (comparisons and adjusted *p* values: Tyr::amiPRT1_ii-Tyr::ami-mCitrine *p*=0.003, all other significant comparisons *p*< 0.001). **d** Two amiRNAs targeted to UBC35 and UBC36 (ami-UBC35/36_i or ami-UBC35/36_ii) or a control (ami-mCitrine). Confocal images were collected one day post-transfection and RFP fluorescence was quantified and plotted as box and whiskers (Empty vector (EV) n= 70, Met::ami-mCitrine n=61, Tyr::ami-mCitrine n=42, Met::ami-UBC35/36_i n=71, Tyr::ami-UBC35/36_i n=84, Met::ami-UBC35/36_ii n=71, Tyr::ami-UBC35/36_ii n=84). Lowercase letters (a, b) denote statistically significant differences between amiRNA constructs, as determined via one-way ANOVA with Tukey’s HSD (comparisons and adjusted *p* values: Tyr::ami-mCitrine-Met::ami-mCitrine *p*=0.02, Met::ami-UBC35/36_i-Tyr::ami-mCitrine *p*=0.001, Met::ami-UBC35/36_ii-Tyr::ami-mCitrine *p*=0.004, Tyr::ami-UBC35/36_ii-Tyr::ami-mCitrine *p*=0.005, all other significant comparisons *p*<0.001). Constructs sharing the same letter are not significantly different.

Using this modified RFP reporter system, we designed two amiRNAs against *At*PRT1 to determine whether this E3 was involved in the endogenous N-degron pathway in *A. thaliana* protoplasts (**Fig 5a, b, c**). Silencing PRT1 in the presence of the Met-CBP-RFP reporter showed no significant difference in the RFP fluorescence signal (**Fig 5b**). Conversely, silencing of PRT1 in the presence of the Tyr-CBP-RFP reporter resulted in an increase in RFP fluorescence compared to the Tyr-CBP-RFP ami-mCitrine control (**Fig 5c**). Confocal microscopy further revealed that PRT1 silencing in protoplasts transformed with Tyr-CBP-RFP resulted in an increase in cytosolic RFP signal (**Supplementary Fig 5**). We also investigated the roles of *At*UBC35 and *At*UBC36 in the N-degron pathway using amiRNAs that were designed to target both functionally redundant genes. Silencing of UBC35 and UBC36 restored the levels of RFP fluorescence in Tyr-CBP-RFP expressing protoplasts to a level comparable to that observed in Met-CBP-RFP expressing protoplasts (**Fig 5d**). This restoration was further evidenced by the localisation of RFP signal within the cytosol in addition to the nuclear localisation inherent in non-silenced Tyr-CBP-RFP constructs (**Supplementary Fig 6**). Collectively, these silencing observations suggest a direct role for PRT1 in destabilising Tyr-CBP-RFP. There are two plausible scenarios that account for the stabilisation of Tyr-CBP-RFP in protoplasts where the levels of UBC35 and UBC36 are knocked down. Scenario one, is that a complex containing at least PRT1 and UBC35 or UBC36 ubiquitinates Tyr-CBP-RFP leading to the degradation of Tyr-CBP-RFP. Scenario two, is that UBC35 or UBC36 ubiquitinate PRT1 leading to a change in localisation or activity of PRT1. These two scenarios do not exclude the possibility that PRT1 interacts with UBC8 family members.

To test the calcium responsiveness of CBP-RFP *in vivo* we transiently expressed Met-CBP-RFP or Tyr-CBP-RFP in *A. thaliana* protoplasts (**Fig 6a**). Protoplasts were exposed to between 0 and 10 mM calcium chloride for 6 hours before imaging. Fluorescence quantified from the Met-CBP-RFP construct was similar at all calcium ion concentrations (**Fig 6b**). Fluorescence from the Tyr-CBP-RFP construct was increased at 2.5 mM calcium relative to the 0 mM control (**Fig 6b**). At higher calcium concentrations there was no significant differences in cytosolic RFP fluorescence, relative to the 0 mM control (**Fig 6b**). These results suggest that addition of exogenous calcium provides a small degree of stabilisation to Tyr-CBP-RFP. This accords with our *in vitro* data (**Fig 3**) where calmodulin binding attenuates, but does not eliminate, PRT1-dependent ubiquitination of Tyr-CBP-RFP.

**Fig. 6.**
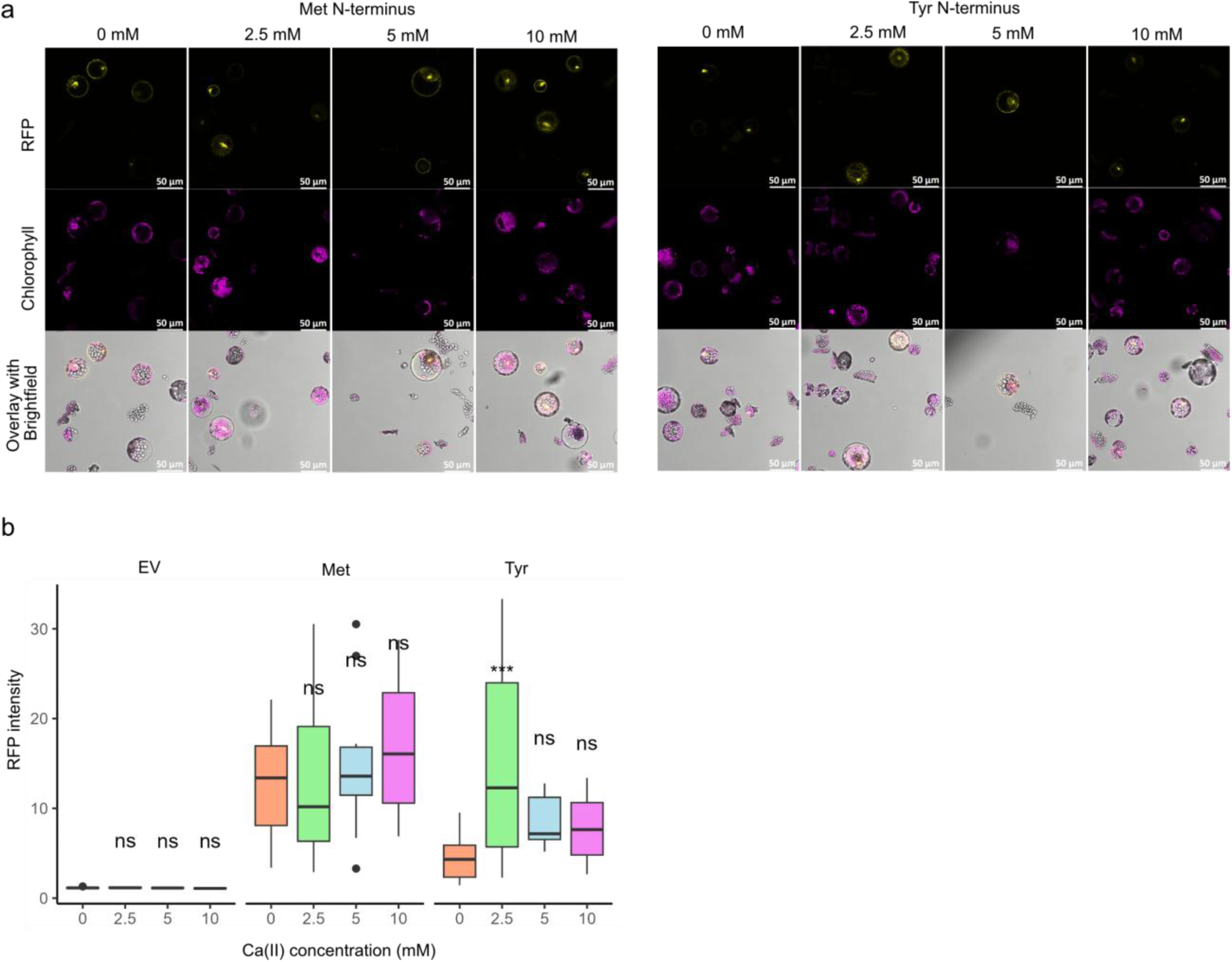
The effect of calcium chloride on CBP-RFP turnover. **a** Representative confocal images of *A. thaliana* protoplasts transiently expressing Met-CBP-RFP, or Tyr-CBP-RFP. Transformed protoplasts were exposed to CaCl_2_ for 6 to 7 hours. RFP fluorescence and chlorophyll autofluorescence is false-coloured yellow and magenta, respectively. Scale bars = 50 µm. **b** Quantification of RFP fluorescence from protoplasts expressing Met-CBP-RFP, Tyr-CBP-RFP or an empty vector control (EV). Fluorescence was quantified in 11-24 positively transformed protoplasts per construct and treatment (EV: 0 mM n=15, 2.5 mM n=16, 5 mM n=11, 10 mM n=17; Met: 0 mM n=16, 2.5 mM n=16, 5 mM n=14, 10 mM n=16; Tyr: 0 mM n=13, 2.5 mM n=24, 5 mM n=12, 10 mM n=15), data were plotted as box and whiskers. Statistical significance was determined via two-way ANOVA with Dunnett’s multiple comparisons test. Not significant (ns) and p<0.001 (***) compared to the 0 mM treatment within each reporter condition.

Having established that RFP can be used as a reporter for PRT1 activity in *A. thaliana* protoplasts, we further probed PRT1 specificity *in vitro*. To separate the reporter protein (RFP) and the recognition element (aromatic N-terminus) into two components, we modified RFP specific LaM-2 nanobodies^34^ so that they have a short N-terminal linker and an aromatic N-terminal residue (**Fig 7a**). All lysine residues in the nanobodies were substituted with arginine to prevent ubiquitination of the nanobody^35^. We produced nanobodies with various N-terminal linkers (between 1 and 9 glycine-serine (GS) repeats) (**supplementary Fig 7**), and as all these nanobodies were functional in our assay we decided to further characterise variants of the most compact nanobody (**Fig 7a**). PRT1 dependent ubiquitination of RFP was observed when nanobodies with Tyr or Trp N-termini were present (**Fig 7b,c**). The apparent rate of RFP ubiquitination was greatest when a nanobody with a Trp N-terminus was present (**Fig 7d**). This is consistent with N-terminal Trp having greater affinity than Tyr for the ZZ domain of PRT1 ^11^.

**Fig 7.**
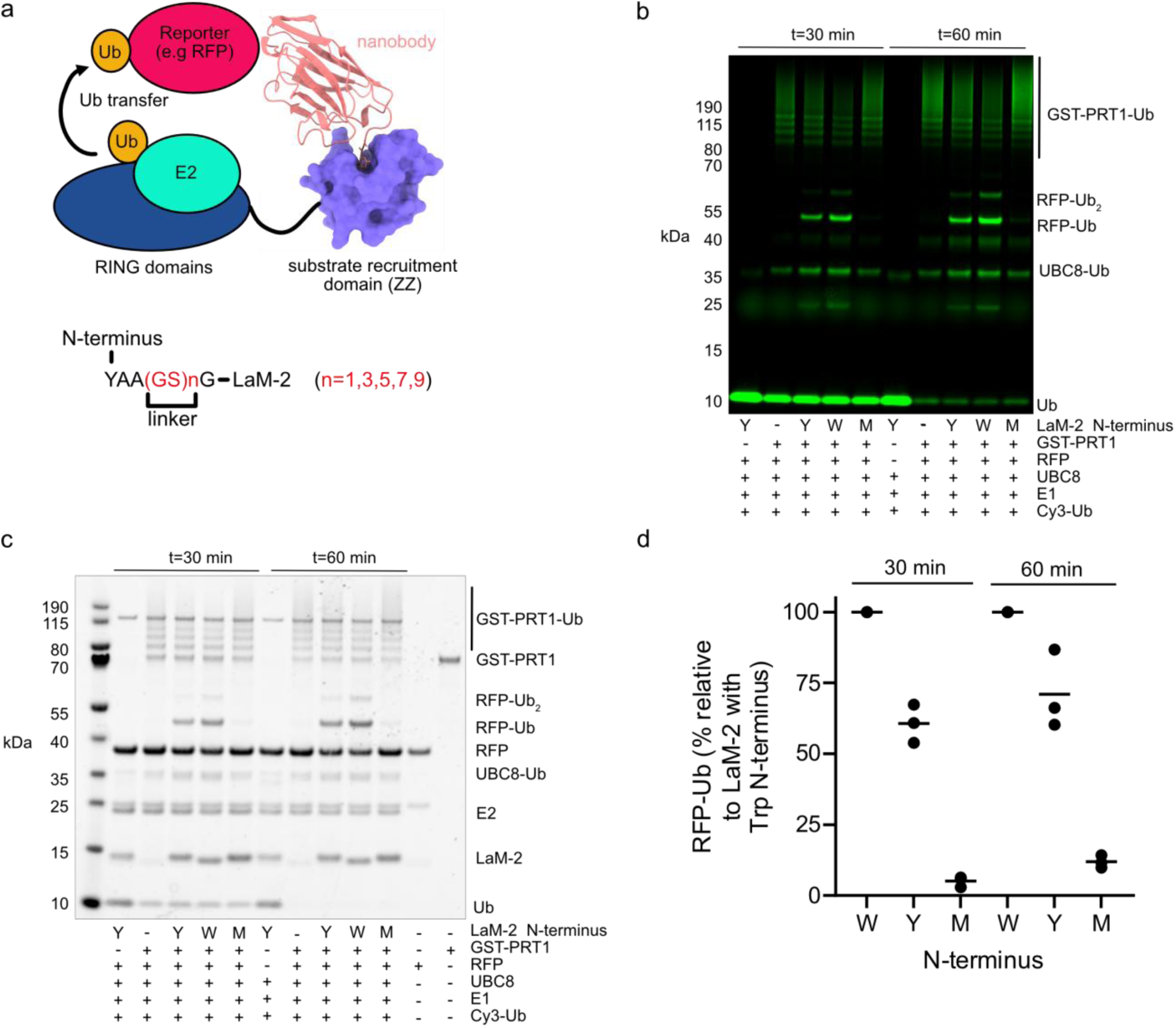
Modified nanobodies acts as substrate adaptors for PRT1. **a** An RFP specific nanobody (LaM-2) with an aromatic amino terminus may be recruited by the ZZ-domain of PRT1 resulting in ubiquitination of RFP. Three-dimensional structures rendered with AlphaFold3. **b** Fluorescently labelled Ub (Cy3-Ub) is transferred to RFP when LaM-2 nanobody with a Tyr (Y) or Trp (W) N-terminus is present (representative gel, n=3). The indicated LaM-2 nanobody, GST-PRT1, RFP, ATP, E1, UBC8, and Cy3-Ub and were incubated for the indicated time. **C** Representative Coomassie blue stained SDS-PAGE gel. **d** Quantification of RFP-Ub in panel b (n=3), bars represent the mean.

The PRT1 protein consists of two RING domains (RING1 and RING2) and a substrate binding ZZ-domain ^11^. The first RING domain of PRT1 has a putative linchpin Arg (PRT1 R66), whereas RING2 does not. In many RING:E2-Ub complexes the linchpin Arg biases the E2-Ub conjugate towards a catalytically productive state, this manifests in enhanced ubiquitination activity ^36–39^. Some RING and RING-like domains lack the linchpin Arg, but they are still competent to bind catalytically productive E2-Ub conjugates ^40,41^. We mutated the linchpin to Ala (PRT1 R66A) and found that this reduced PRT1 dependent Ub transfer from both UBC8 and UBC35 (**Fig 8a, b, c**). This suggests that both UBC8 and UBC35 are activated by RING1.

**Fig 8.**
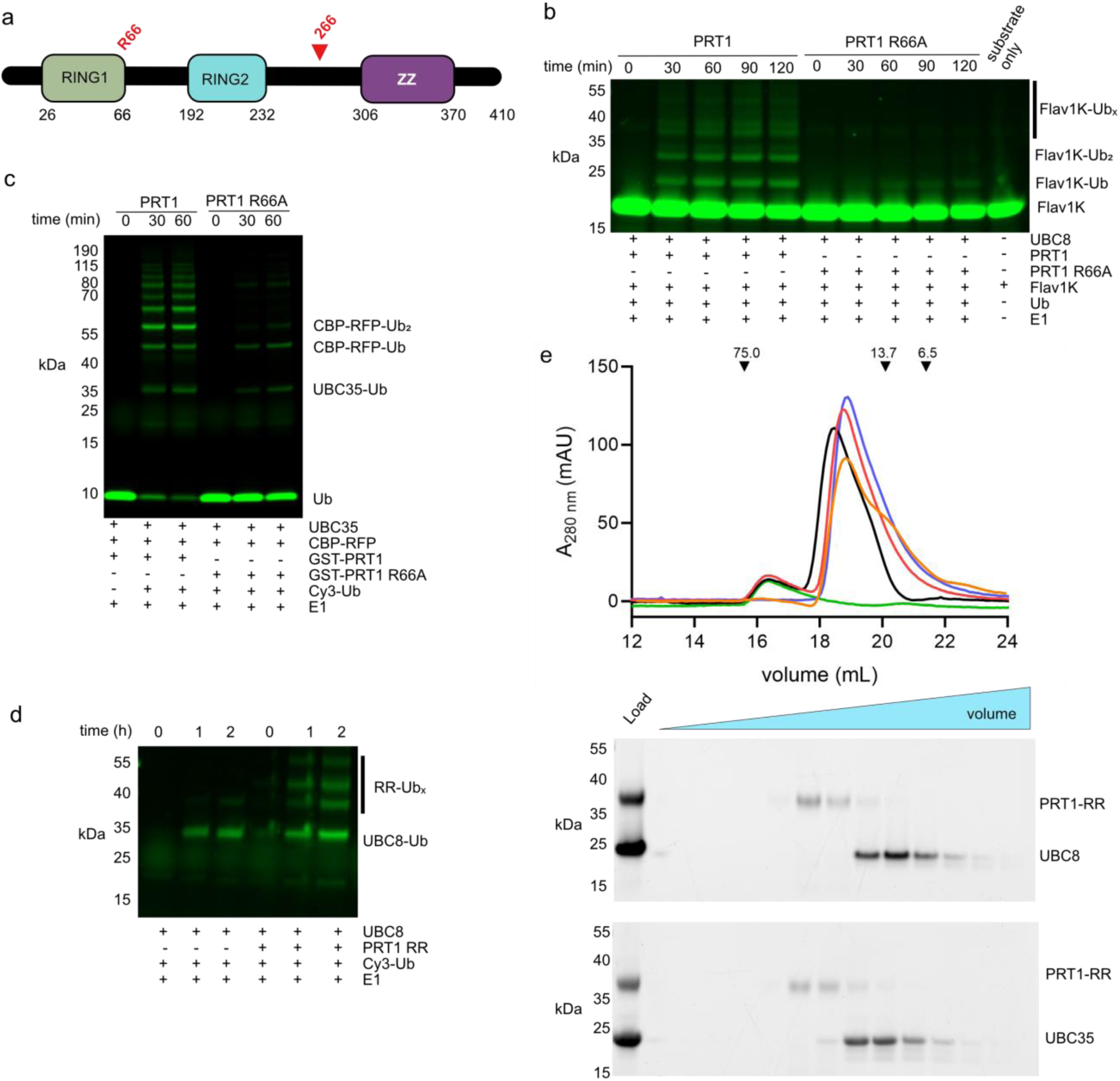
Both UBC8 and UBC35 bind to RING1 of PRT1. **a** Schematic of PRT1 indicating the position of the putative linchpin Arg (R66). The protein was truncated after position 266 to create PRT1-RR domain. **b** Mutation of the linchpin Arg to Ala results in reduced activation of UBC8, representative gel with fluorescent substrate (n=2). GST-PRT1 or GST-PRT1 R66A, FlaV1K, ATP, E1, UBC8, and Ub were incubated for the indicated time. **c** The linchpin Arg is required for transfer of Ub from UBC35 to a CBP-RFP (representative gel, n=2). GST-PRT1 or GST-PRT1 R66A, CBP-RFP, ATP, E1, UBC35, and Ub and were incubated for the indicated time. **d** The PRT1-RR domain (2 µM) activates UBC8 (2.5 µM). PRT1-RR domain, UBC8, ATP, E1 and Cy3-Ub were incubated for the indicated time. **e**. Size-exclusion chromatography (SEC; Superdex S200 Increase 10/300 GL) chromatograms of PRT1-RR (green), UBC8 (blue), UBC35 (orange), PRT1-RR+UBC8 (red), PRT1-RR+UBC35 (black). Proteins (5 µM PRT1-RR, 20 µM UBC8, or UBC35) were incubate on ice for 1 hr prior to SEC. The elution volumes and molecular masses of standards are indicated with arrows. Coomassie blue stained gels of the eluted proteins are shown below the chromatogram.

To explore the affinity of E2s for PRT1, we conducted size-exclusion chromatography (SEC) of the RING1-RING2 domain (Residues 1-273, henceforth PRT1-RR), and UBC8, or UBC35. The PRT1-RR domain had intrinsic activity (**Fig 8 d**); however, we did not observe stable complex-formation between PRT1-RR and UBC8 or UBC35 (**Fig 8e**). This is consistent with the modest affinity of isolated RING domains for E2s ^42,43^. For some N-recognins (UBR1, PRT6, UBR4, and BIG), auxiliary non-RING regions contribute to E2 binding^7,9,40,41,44,45^, but there are no homologous non-RING regions in PRT1. Taken together our data suggest a model where PRT1 interacts with substrates, including E3 ligases that are themselves active with UBC8 family members (**Fig 9a**). This PRT1-substrate interaction facilitates the recruitment of UBC35-UEV1A to the substrate (**Fig 9b**), or the recruitment of a UBC8 family member (**Fig 9c**). Both UBC8 and UBC35 are likely to occupy a single binding site (PRT1 RING1), but there is nothing to prevent the E2s from acting sequentially.

**Fig 9.**
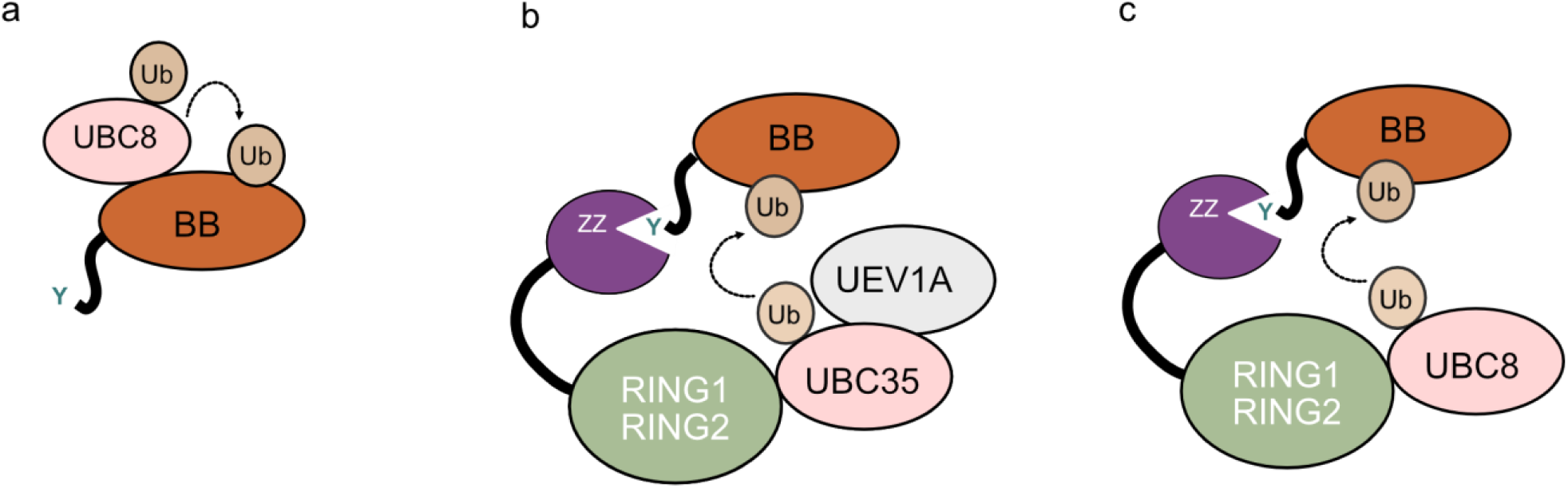
Model of interactions between PRT1, UBC8, UBC35-UEV1A and a substrate. **a** A proteolytic fragment of BIG BROTHER with an N-terminal Tyr (Y61 BB) is an active E3 ligase, which is ubiquitylated by UBC8 family members. **b** PRT1 has two RING domains (RING1 and RING2) and a substrate binding ZZ domain. The ZZ domain of PRT1 binds to the N-terminus of Y61 BB. A residue within RING1 of PRT1 (Arg66) is required for activation of both UBC35 and UBC8. Both the PRT1-UBC35 and PRT1-UBC35-UEV1A complexes transfer Ub to Y61-BB. **c** UBC8 family members transfer Ub to Y61-BB in both PRT1-dependent and PRT1-independent reactions.

## Conclusions

The PRT1 and PRT6 proteins have affinity for aromatic N-degrons, and it has been assumed that *in planta* both E3s build pro-degrative K48-linked Ub chains ^8,11,13^. Our *in vitro* data suggests that PRT1 interacts with UBC35-UEV1A to build K63-linked Ub chains. It is possible that *in vivo* PRT1 interacts with UBC35-UEV1A to build K63-linked Ub chains on substrates. As K63-linked Ub chains are associated with autophagy^46^ this observation may offer a new avenue to explore the phenotypes of *prt1* mutant plants^3,4^. In Arabidopsis UEV1A is anchored to membranes^25^, and it is possible that PRT1 also directs ubiquitination of membrane associated N-degrons. However, the only known substrate of PRT1 is a proteolytic fragment of BIG BROTHER (Y61-BB). The Y61-BB protein is itself an E3 ligase that is active with UBC8 family members^20,28^. It has been difficult to reconcile the high, UBC8 dependent, autoubiquitination activity of Y61-BB with the apparent PRT1-UBC8 dependent ubiquitination of Y61-BB^11^. We found that Y61-BB was ubiquitinated by UBC35 and the UBC35-UEV1A complex only when PRT1 was present. Our *in vitro* data do not allow us to make conclusions about the physiological significance of the PRT1-UBC35-UEV1A interaction with Y61-BB, but our data does suggest the interaction could occur if the proteins are collocated in the cell.

Across our experiments PRT1-UBC8 was more active than PRT1-UBC35-UEV1A with all the substrates we analysed in our *in vitro* assays. This may reflect the intrinsic high-activity of UBC8^22^, as well as the ability of the UBC35-UEV1A complex to build K63-linked Ub chains^27,47^. The observed inactivity of UBC35-UEV1A with a single lysine substrate (Flav1K) may be due to the reaction stoichiometry. In this reaction, the amount of ubiquitin available for K63 chain formation exceeded the amount of PRT1-bound substrate.

To explore the roles of PRT1 and UBC35 in substrate degradation we developed a system to transiently express a reporter protein and transiently knockdown components of the protein degradation-machinery. We analysed transient expression of fluorescent reporter proteins with stabilising (Met) or destabilising (Tyr) N-termini in *A. thaliana* protoplasts and *N. benthamiana* leaves. The fluorescence signal from the destabilised construct was two to three times less than that from the stable construct. We were able to analyse the effects of knocking down PRT1 or UBC35 and UBC36 on Tyr-CBP-RFP stability, by placing the respective amiRNA within the CBP-RFP gene. We found that the stability of Tyr-CBP-RFP increased when PRT1 or UBC35 and UBC36 were knocked down. Increased stability of Tyr-CBP-RFP when PRT1 was knocked down, supports the view that PRT1 is the primary plant N-recognin for aromatic N-degrons ^11,15^. The roles of UBC35 and UBC36 in destabilising Tyr-CBP-RFP are less clear. Others have shown that knocking out UBC35 and UBC36 has severe pleotropic effects in *A. thaliana* ^26,27^, and we cannot rule out global changes to proteostasis in our UBC35 and UBC36 knockdown experiments. One possibility, supported by our data, is that a complex consisting of at least PRT1 and UBC35 or UBC36 is responsible for the degradation of Tyr-CBP-RFP. This possibility does not rule out interactions between UBC8 family members and PRT1.

In mammals the autophagy complex p62/Sequestosome-1 is an attractive candidate for targeted protein degradation (TPD) ^48–50^. Similarly, the PRT1, PRT6 and BIG E3 ligases could be exploited for TPD in plants^16^. Our nanobodies provide a simple platform for screening the interactions of PRT1 with substrates. A caveat is that the N-termini of nanobodies are likely to be recognised by BIG and PRT6 as well as PRT1 ^8,51^. There is also the possibility that the nanobodies will act as competitive inhibitors of N-recognins ^52^. Interestingly, the nanobodies with a single GS repeat behave in a similar manner to the nanobody with 9 GS repeats under our assay conditions. This suggests that many regulatory motifs could be accommodated at the N-terminus of the nanobody. In principle a cellular protease (rather than TEV) induced upon specific stimuli, could be used to cleave the nanobody to reveal the aromatic residue and recruit an N-recognin.

To our knowledge, we are the first to report a synthetic calmodulin-controlled off-switch for ubiquitination. In principle, such a system could be fine-tuned to trigger TPD in response to physiologically relevant changes in calcium ion concentration^53^. In many plant species exogenously applied calcium results in increases in intracellular calcium concentration^54^. There is evidence that calmodulin interacts with naturally occurring E3 ligases. For example, both mammalian UBR4 and *A. thaliana* BIG form tight complexes with calmodulin ^9,44,55–57^, and both E3s are associated with calcium signalling ^56,58^. Our *A. thaliana* protoplast data show there was a modest stabilisation of the reporter (Tyr-CBP-RFP) when protoplasts were exposed to elevated calcium. The utility of this reporter is limited by its dynamic range and protoplast to protoplast variability in transgene expression. As such, the present design would not be expected to respond to the transient spikes in calcium ion concentration which have been observed in plants^53^. We expect that our nanobodies will be broadly useful in delineating N-degron pathways across biology, and that our calmodulin regulated-constructs will be valuable baselines in developing calcium controlled TPD tools.

## Methods

### Protein expression constructs

Complete amino acid sequences for constructs produced in this study are provided in the supplementary methods. Codon optimised sequences encoding UEV1A (At1g23260). UEV1C (At2g36060), UBC1 (At1g14400), UBC4 (At5g41340), UBC7(At5g59300), UBC8 (At5g41700), UBC22 (At5g05080), UBC27 (At5g50870), and UBC35 (At1g78870) were cloned into pET-28a(+) expression vectors by Twist Bioscience. The resulting proteins have six-histidine tags at both termini. A codon optimised sequence encoding residues of *A. thaliana* PRT1 (At3g24800) was cloned into a pGEX-6P-1 vector by GenScript. The resulting protein has an N-terminal glutathione s-transferase (GST) tag followed by a 3C-protease cleavage site. Vector encoding UBE1 and Ub have been described elsewhere^59,60^.

A pET-28a(+) expression vector encoding Ub with an N-terminal 3C-protease site and Cys-Gly linker was produced by Twist Bioscience. Following 3C-protease cleavage the resulting Ub has Gly-Pro-Cys-Gly at the N-terminus. Labelling of the single cysteine, of Ub or Flav1K, with Cy3 maleimide was achieved as described previously^61^. A pET-28a(+) expression vector encoding Flavodoxin (PDB 2M6R^62^) with three Lys to Arg substitutions (K32R, K76R, K105R), an N-terminal six-histidine tag and a TEV proteases cleave site, was produced by Twist Bioscience. A gene encoding the red fluorescence protein (RFP) variant mCherry ^29^ was cloned into a pET-28a(+) expression vector by Twist Bioscience. Similarly, a pET-28a(+) expression vector encoding *A. thaliana* BIG BROTHER (At3g63530) residues 61-248, with an N-terminal six-histidine tag and TEV-protease cleavage site and a C-terminal FLAG epitope was produced by Twist Bioscience. A gene encoding *A. thaliana* CaM4 (At1g66410) with an N-terminal 3C-protease site was cloned into a pET-28a(+) expression vector by Twist Bioscience.

LaM-2 nanobody residues 3-127 with an N-terminal tobacco etch virus protease (TEV) site (Asn-Leu-Tyr-Phe-Gln-Xaa, where Xaa is any amino acid) and linker sequences were cloned into pET-28a(+) expression vectors. The resulting proteins have an N-terminal six-histidine tag. Cleavage of the nanobody by TEV protease results in Xaa at the N-terminus. A gene encoding CBP-RFP with an N-terminal six histidine tag and TEV protease cleavage site was synthesised by IDT and ligated into the pET-28a(+) expression vector.

### Protein expression

All constructs were expressed in *E. coli* BL21(DE3) cells cultured in LB media. Cells were grown at 37 °C until they reached an optical density (600 nm) of 0.6. At this point, cells were chilled to 16 °C and protein expression was induced for 16 hours with 0.3 mM IPTG.

### Protein purification

Ubiquitin was purified using a protocol modified from ^59^. Cells were resuspended in 50 mM ammonium acetate (pH 4.5) and lysed by sonication. Clarified lysate was applied to a SP-Sepharose Fast Flow column (Cytiva) equilibrated in 50 mM ammonium acetate (pH 4.5). Ubiquitin was eluted in 50 mM ammonium acetate (pH 4.5) using a linear gradient of 0–500 mM NaCl. Fractions containing Ub were pooled and exchanged, by size-exclusion-chromatography (SEC) (Superdex S75, Cytiva), into 20 mM HEPES-NaOH (pH 7.5), 150 mM NaCl, 1 mM TCEP (tris(2-carboxyethyl)phosphine). Six-histidine and GST tagged proteins were purified using standard techniques. Proteolytic digestions with TEV or 3C protease, were performed overnight at 4**°**C. Cleaved proteins were passed back through the appropriate affinity resin prior to SEC. All proteins were exchanged by SEC (Superdex S75 or Superdex S200, Cytiva) into 20 mM HEPES-NaOH (pH 7.5), 150 mM NaCl, 1 mM TCEP. Protein aliquots were snap frozen in liquid nitrogen and stored at -70 **°**C.

### Fluorescent Ub discharge assays

Assays were carried out at 30 **°**C in assay buffer (20 mM HEPES-NaOH (pH 7.5), 150 mM NaCl, 0.5 mM TCEP, 5 mM ATP, 5 mM MgCl_2_) with 3 µM Cy3-Ub, 0.3 to 0.7 µM UBE1, 1 uM GST-PRT1, 2.5 µM UBC8, UBC35, UEV1A, or UEV1C, 5 µM nanobody, 20 µM AtCaM4, 5µM RFP, or 5 µM CBP-RFP. Assays were stopped by the addition of lithium dodecyl sulfate (LDS) sample buffer (ThermoFisher) and 5 % (v/v) 2-mercaptoethanol. Samples were electrophoresed in NuPAGE Bis-Tris Mini Protein Gels, 4–12% gels (ThermoFisher). Following SDS-PAGE gels were imaged with a Chemidoc Gel Imaging System (BioRad) or an iBright Imaging System (ThermoFisher). Quantification was achieved using ImageJ v 1.53 and data were graphed using GraphPad Prism (Graphpad Software).

### Immunoblot

Following SDS-PAGE samples were transferred to nitrocellulose membranes using the Trans-Blot Turbo system (Bio-Rad), membranes were blocked and immunoblotted with the DYKDDDDK Tag Monoclonal Antibody (FG4R), DyLight 680 (ThermoFisher). Membranes were imaged using a Chemidoc Gel Imaging System (BioRad).

### Ubiquitin discharge assays

Assays were carried out at 30 **°**C in assay buffer with 50 µM Ub, 0.3 to 0.7 µM UBE1, 1 µM GST-PRT1, 2.5 µM UBC8, UBC35, UEV1A, UEV1C, 3.4 µM Flav1K, or 5 µM CBP-RFP. Assays were stopped and visualised as described for Fluorescent Ub discharge assays.

### Transient expression vectors

Plasmid constructs were assembled using Golden Gate (Type IIS) MoClo cloning strategies^63^. Ubiquitin leader sequences, CBP domains with variable N-termini (Met and Ala) and the mCherry variant of RPF modified with intron splice sites were synthesised by Integrated DNA Technologies (IDT). The amiRNA sequences for *At*PRT1 (At3g24800), *At*UBC35 (At1g78870) and *At*UBC36 (At1g16890) were designed using WMD3 Web MicroRNA Designer (http://wmd3.weigelworld.org/cgi-bin/webapp.cgi) and based on the Arabidopsis miR319a scaffold. These pre-amiRNA sequences were also synthesised by IDT for cloning. A dummy sequence (5’-TATACGTAA-3’) without an amiRNA scaffold was used to provide the necessary overhangs for correct assembly of the Met-CBP-RFP:Dummy construct. All constructs were cloned into the F2 acceptor vector (pICH47742)^64^, with expression driven by the 35S Cauliflower Mosaic Virus promoter + Tobacco Mosaic Virus omega (5’ UTR) sequence (pICH51266)^63^ and terminated by the AtHSP18 terminator sequence.

### Plant growth conditions

Wildtype (WT) *A. thaliana* Col-0 plants were stratified for 48 hr and germinated on seed raising mix supplemented with 3 g/L Osmocote Exact Mini in 400 mL punnets. Plants were grown in a controlled temperature room under a 16 hr light/8 hr dark photoperiod, with temperature maintained at 21 °C, a relative humidity level of 55% and light intensity of 110 μmol photons/m^2^/s. Only healthy, fully expanded rosette leaves were used for protoplast isolation.

WT *N. benthamiana* seeds were sown in individual 135 mm (1.5 L) round plastic pots containing seed raising mix supplemented with Osmocote Exact Mini, as described above. Plants were germinated and grown in a controlled temperature room as described for *A. thaliana,* except with a higher light intensity of 325 μmol photons/m^2^/s. Only the first, second and third true leaves from 4-5 week-old *N. benthamiana* plants were used for agroinfiltration, while the rest were discarded.

### Protoplast isolation and transient transfection

Protoplasts were isolated from 5-6 week-old *A. thaliana* rosette leaves, following established protocols^65,66^, with the following modifications. Fully expanded rosette leaves were prepared for digestion by removing the abaxial epidermis via the Tape Sandwich method. One rosette leaf was digested per mL of Digestion Buffer (DB; 20 mM MES pH 5.7, 0.4 M sorbitol, 10 mM KCl, 1.2% (w/v) Cellulase RS, 0.6% (w/v) Macerozyme R-10, 10 mM CaCl_2_, 0.1% (w/v) BSA) for 30 min at room temperature (RT) with shaking at 50 rpm. An equal volume of cold Wash Buffer (WB; DB without Cellulase RS and Macerozyme R-10) was added to stop digestion. Crude protoplasts were collected in round bottom microfuge tubes by centrifuging in a swing bucket rotor at 100 g for 3 min. Crude protoplast pellets were resuspended in Dex10 (Suc20 supplemented with 10% (w/v) dextran T40), layered with Suc20 (20 mM MES pH 5.7, 20% (w/v) sucrose, 10 mM KCl, 10 mM CaCl_2_) and WB to form a density gradient. The density gradient was centrifuged at 100 g for 5 min, after which purified protoplasts were collected from the upper interphase and washed with WB. Purified protoplasts were pelleted as before and resuspended in SMg (4 mM MES pH 5.7, 0.4 M sorbitol, 15 mM MgCl_2_). Protoplast viability and concentration were assessed on a hemocytometer with 0.2% (v/v) Trypan Blue.

For each independent transformation, purified plasmid DNA and protoplasts were mixed in a transfection ratio of 5 µg DNA: 5000 protoplasts. PEG-40 (40% (w/v) PEG-4000, 0.2 M sorbitol, 100 mM CaCl_2_) was incubated with each reaction for 5 min to facilitate transformation before terminating with 2X volume of W5 (2 mM MES pH 5.7, 154 mM NaCl, 125 mM CaCl_2_, 5 mM KCl). Transfected protoplast reactions were centrifuged at 100 g for 3 min in a swing bucket rotor and resuspended in 250 µL WI (4 mM MES pH 5.7, 0.4 M sorbitol, 20 mM KCl) to facilitate ease of transfer into flat bottom plates for incubation. Plates were incubated at RT with constant shaking at 50 rpm for at least 16 hr to enable the expression of plasmid constructs. Protoplasts transfected with amiRNA constructs were incubated for at least 24 hr to enable sufficient silencing.

### Addition of calcium to protoplasts

Transformed *A. thaliana* protoplasts were treated with CaCl_2_ at 1-day post transfection to final concentrations of 0, 2.5, 5 and 10 mM. A concentrated 500 mM stock solution of CaCl_2_ was used to prevent significantly diluting the sorbitol in the WI solution and bursting fragile protoplasts. Protoplasts were left to incubate at RT and 50 rpm shaking for 6-7 hr prior to imaging via confocal microscopy.

### Agroinfiltration of *N. benthamiana* leaves

Reporter constructs were transiently expressed in *N. benthamiana*, following established protocols^67,68^. Briefly, *Agrobacterium tumefaciens* strain GV3101 (pMP90) transformed with the constructs of interest were cultured in LB media with 25 µg/mL rifampicin, 50 µg/mL gentamycin and 50 µg/mL carbenicillin at 28 °C and 220 rpm for 24 hr. *A. tumefaciens* strain GV3101 (pMP90) containing a plasmid encoding the Tomato Bushy Stunt Virus P19 viral suppressor protein^69^ was also mixed with cultures containing the plasmid(s) of interest at optical densities (OD 600 nm) of 0.3 and 0.8, respectively for co-infiltration. A P19-only control was prepared to an OD_600_ of 0.8 as a negative control. Following overnight growth, bacterial cultures were pelleted at 2,150 g for 8 min and resuspended in infiltration solution (10 mM MES pH 5.7, 10 mM MgCl2, 150 μM acetosyringone). Resuspended cells were incubated at RT for 2 hr prior to infiltration into the abaxial surface of 4-5 week-old *N. benthamiana* leaves using a needle-less syringe. Transient protein expression was assessed 3 days post-infiltration (dpi) via fluorescence screening on a ChemiDoc™MP Imaging System (Bio-Rad) using green epi-illumination coupled with a 605/50 filter. High-resolution images were captured Leica TCS SP8 confocal microscope.

### Confocal microscopy

Confocal laser scanning microscopy was performed on transiently infiltrated *N. benthamiana* leaves and transfected *A. thaliana* protoplasts expressing the various CBP-RFP constructs of interest. Images were taken on a Leica SP8 confocal laser microscope using a ×20 water immersion objective, PMT detectors and the Leica Application Suite X (LAS X) software package. mCherry fluorescence and chlorophyll fluorescence was captured via sequential scans between frames to eliminate spectral bleed-through. The following excitation and emission wavelengths were used to detect the respective fluorophores in *N. benthamiana*: mCitrine at λex = 514 nm and λem = 525-555 nm, mCherry at λex = 561 nm and λem = 590-620 nm, and chlorophyll at λex = 514 nm and λem = 630-690 nm. The following excitation and emission wavelengths were used to detect the respective fluorophores in *A. thaliana*: mCherry at λex = 561 nm and λem = 590-620 nm, and chlorophyll at λex = 561 nm and λem = 650-690 nm. All image processing post-capture was performed in LAS X Office (version 1.4.5).

### Data visualisation and statistical analysis

mCherry fluorescence captured via confocal microscopy in successfully transfected *A. thaliana* protoplasts was quantified using FIJI^70^ to account for variable protoplast transformation efficiencies across batches. mCherry fluorescence screened in transiently infiltrated *N. benthamiana* leaves on the ChemiDoc™MP Imaging System (BioRad) was quantified using the volume tool in Image Lab (version 6.0, BioRad). Data visualisation, processing, and statistical analyses were performed in R (version 4.3.1) with R Studio and the ggplot2 package^71^. To compare *in vivo* mCherry fluorescence across constructs under steady-state and silencing conditions, a one-way ANOVA followed by Tukey’s multiple comparisons test was performed. To assess the effect of various CaCl_2_ concentrations on mCherry fluorescence in protoplasts, a two-way ANOVA with Dunnett’s multiple comparisons test was performed.

**Table 1:**
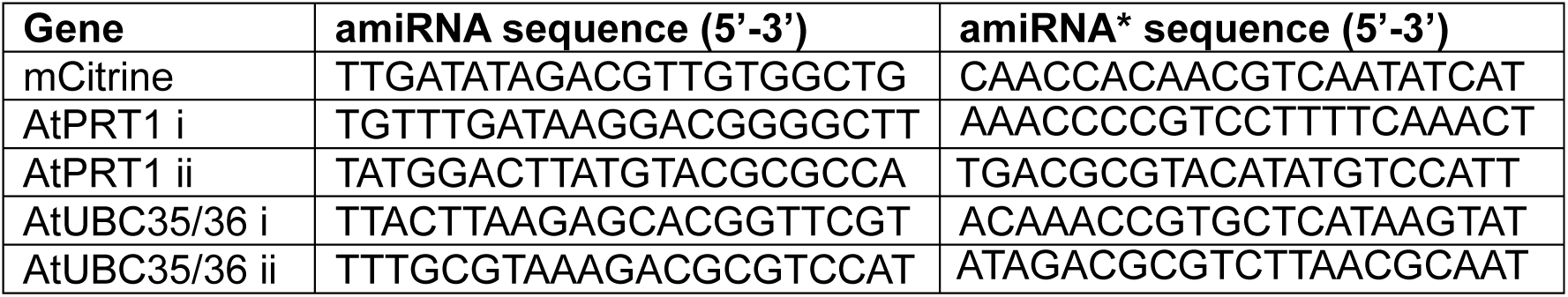
amiRNA sequences designed to target components of the protein degradation machinery in *A. thaliana*. The active guide strand of the mature amiRNA duplex is designated as amiRNA, while its complementary passenger strand is shown with a star (*) symbol.

## Contributions

P.D.M. conceived the research and obtained funding. K.E.A.O. & P.D.M. designed and carried out *in vitro* experiments. S.Y. & K.X.C. designed and carried out *in vivo* experiments. P.D.M. wrote the manuscript with assistance from K.E.A.O., S.Y., & K.X.C

## Acknowledgments

We thank Catherine Day and Claudia Rossig (University of Otago) for the kind gift of plasmids encoding Ub and E1. We thank Adam Fletcher (University of Glasgow) for critically reading the manuscript.

