## Supplementary data for "Synthetic tools to redirect the ubiquitin E3 ligase activity of, PRT1, a plant-specific N-recognin"

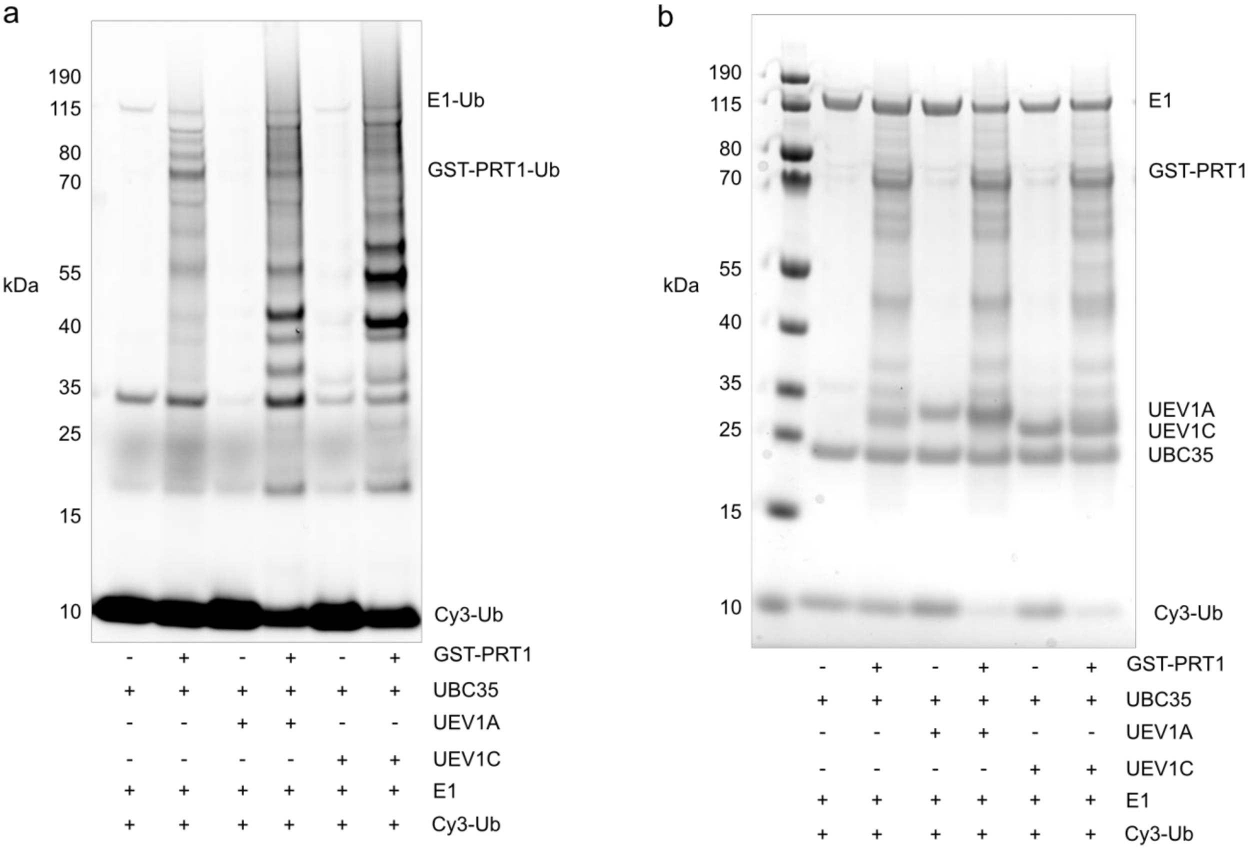


**Supplementary Fig 1. Transfer of Ub from UBC35, UBC35-UEV1A or UBC35-UEV1C to PRT1. a** GST-PRT1, UBC35, UEV1A, UEV1C, E1 and Cy3-Ub were incubated for three hours. Samples were resolved by SDS-PAGE and Cy3 fluorescence was imaged. **b** Coomassie blue stained gel**.**

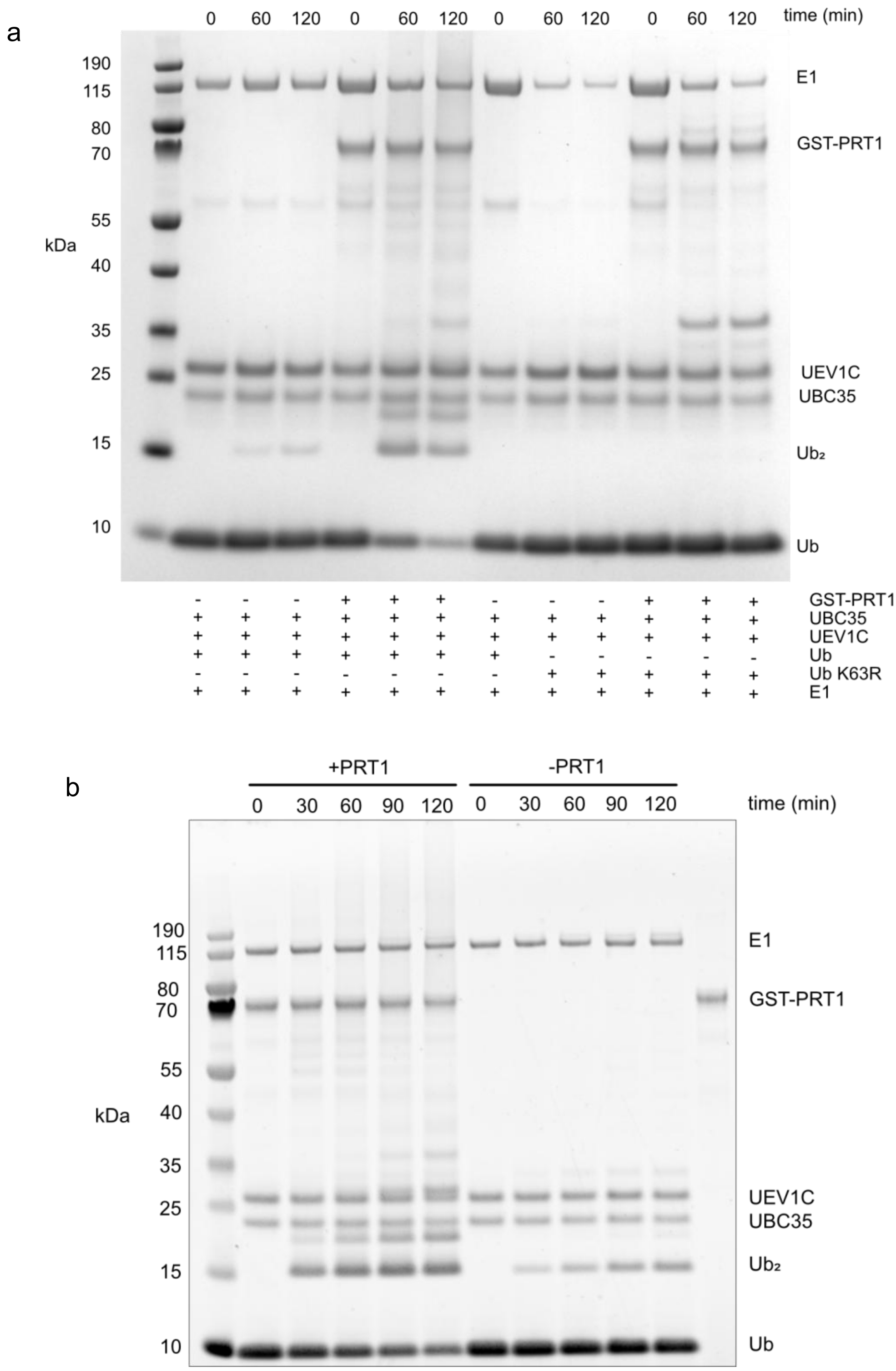


**Supplementary Fig 2. PRT1 interacts with the UBC35-UEV1C complex. a** GST-PRT1, UBC35, UEV1C, E1 and Ub or Ub K63 were incubated for the indicated times. **b** GST-PRT1, UBC35, UEV1C, E1 and Ub were incubated for the indicated times. Samples were resolved by SDS-PAGE, and the gels were stained with Coomassie blue**.**

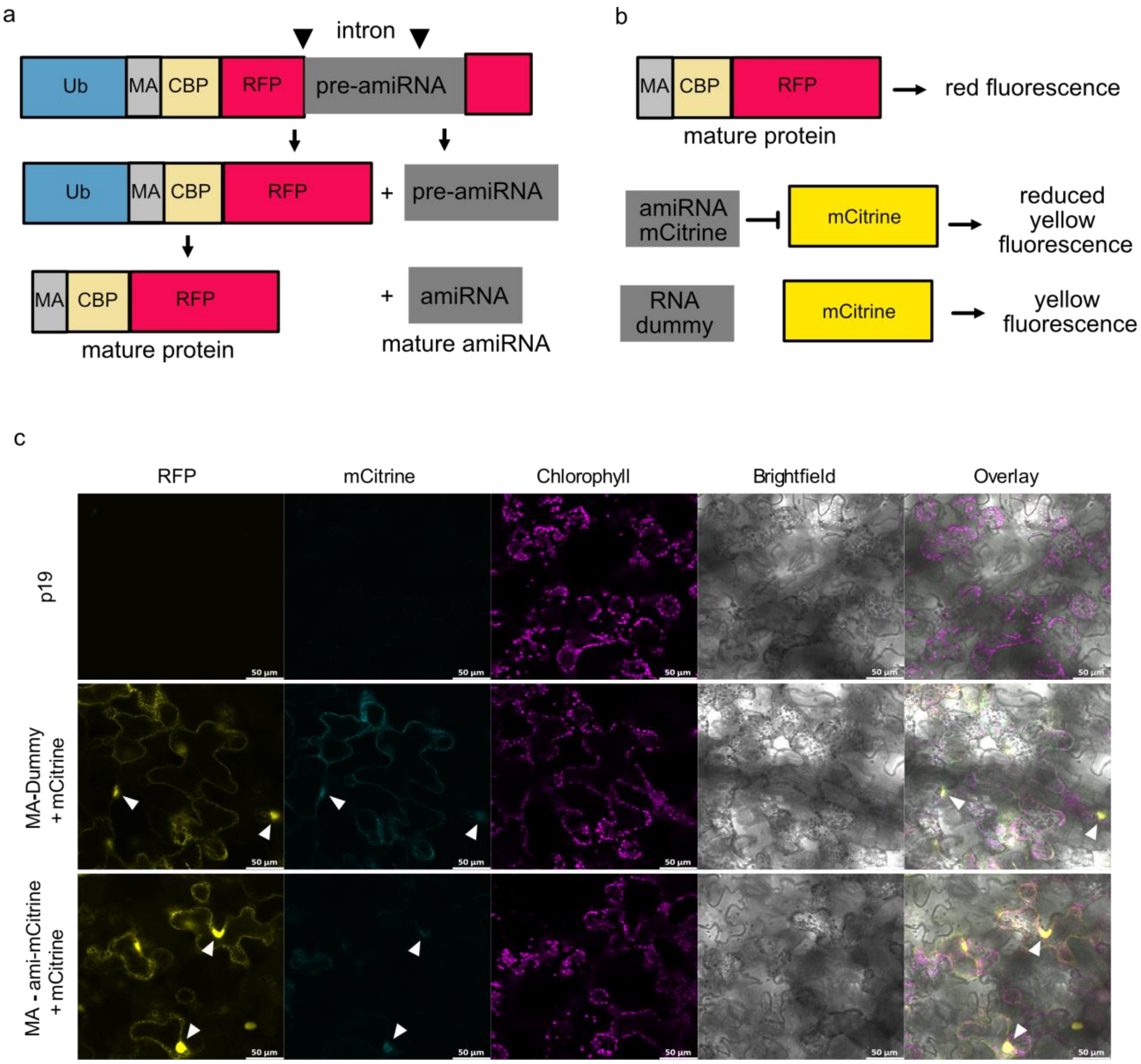


**Supplementary Fig 3. Evidence of artificial microRNA (amiRNA) splicing and subsequent mCitrine silencing in *N. benthamiana* leaves**. **a** Schematic of Ub-CBP-RFP fusion with a pre-amiRNA present as an intron. Correct splicing of the two exons is required for the translation of the fluorescent reporter protein. **b** The CBP-RFP protein with a stabilising (Met) N-terminus produces red fluorescence. An amiRNA targeting mCitrine reduces expression of mCitrine and consequently reduces the level of yellow fluorescence. The RNA dummy is DNA sequence (without the pre-amiRNA scaffold) cloned into the intron of mCherry **c** The RNA Dummy (control) or ami-mCitrine sequences were introduced as introns within the CBP-RFP reporter. The two CBP-RFP reporter constructs were transiently co-expressed with mCitrine in *N. benthamiana* leaves. Representative confocal images of Dummy, ami-mCitrine and p19 (negative control) infiltrated leaves. All leaves were co-infiltrated with *Agrobacterium tumefaciens* strain GV3101 (pMP90) containing a plasmid encoding the Tomato Bushy Stunt Virus P19 viral suppressor protein. Images were collected at 3 days post infiltration. RFP fluorescence is false-coloured yellow, while mCitrine fluorescence is false coloured cyan and chlorophyll autofluorescence is denoted in magenta. White arrows indicate nuclei within transformed leaf mesophyll cells. Scale bars = 50 µm.

a

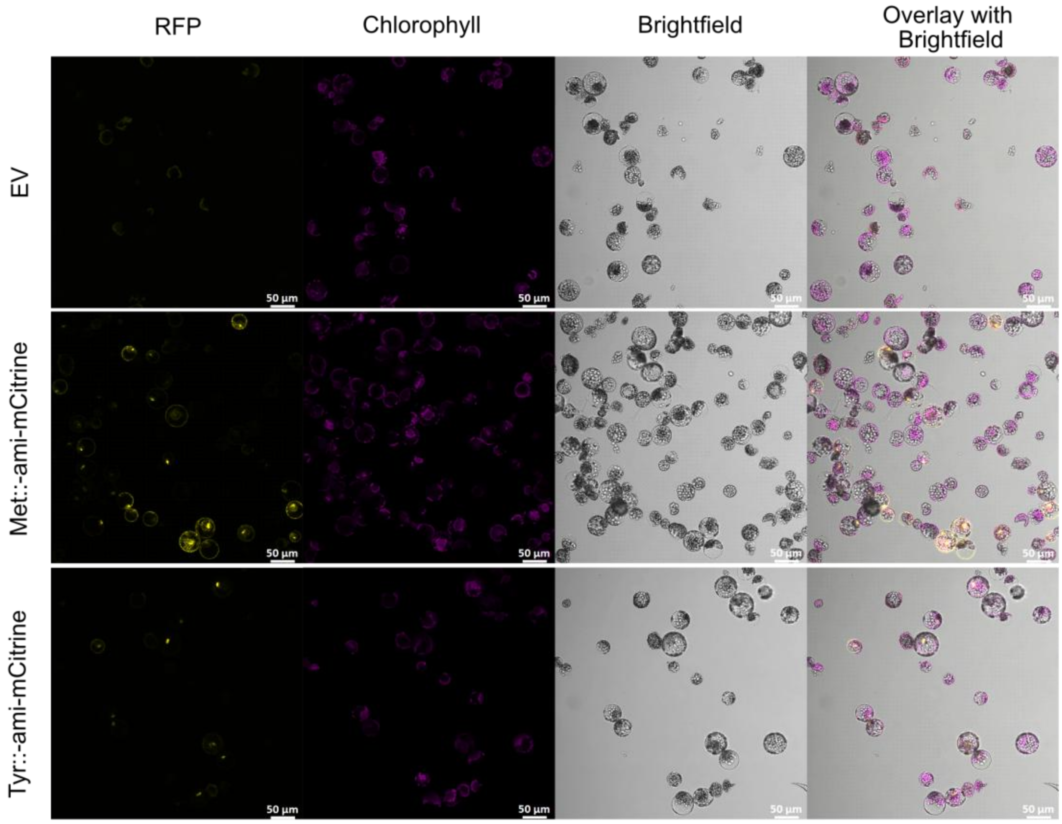


b

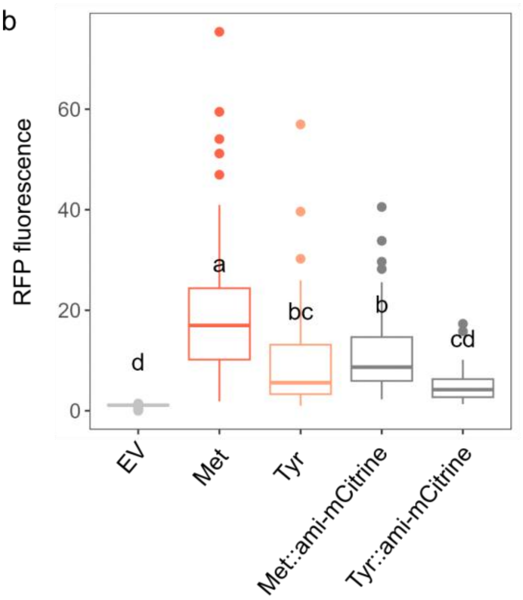


**Supplementary Fig 4. Expression of CBP-RFP with ami-mCitrine present as an intron within the CBP-RFP gene. a** Representative confocal images of *A. thaliana* protoplasts expressing Met-CBP-RFP and ami-mCitrine (Met::ami-mCitrine), Tyr-CBP-RFP and ami-mCitrine (Tyr::ami-mCitrine) or empty vector (EV). Images were collected at one day post-transfection. RFP fluorescence is false-coloured yellow, chlorophyll autofluorescence is denoted in magenta. Scale bars = 50 µm. **b** RFP fluorescence quantified in 42-117 positively transformed *A. thaliana* protoplasts per construct and plotted as box and whiskers (EV n=70, Met-CBP-RFP n=117, Tyr-CBP-RFP n=59, Met::ami-mCitrine n=61, Tyr::ami-mCitrine n=42). Lowercase letters (a, b) denote statistically significant differences from a one-way ANOVA with Tukey's HSD test. Constructs sharing the same letter are not significantly different. (comparisons and adjusted $p$ values: Tyr::ami-mCitrine- Met::ami-mCitrine $p$=0.006, all other significant comparisons $p$<0.001)

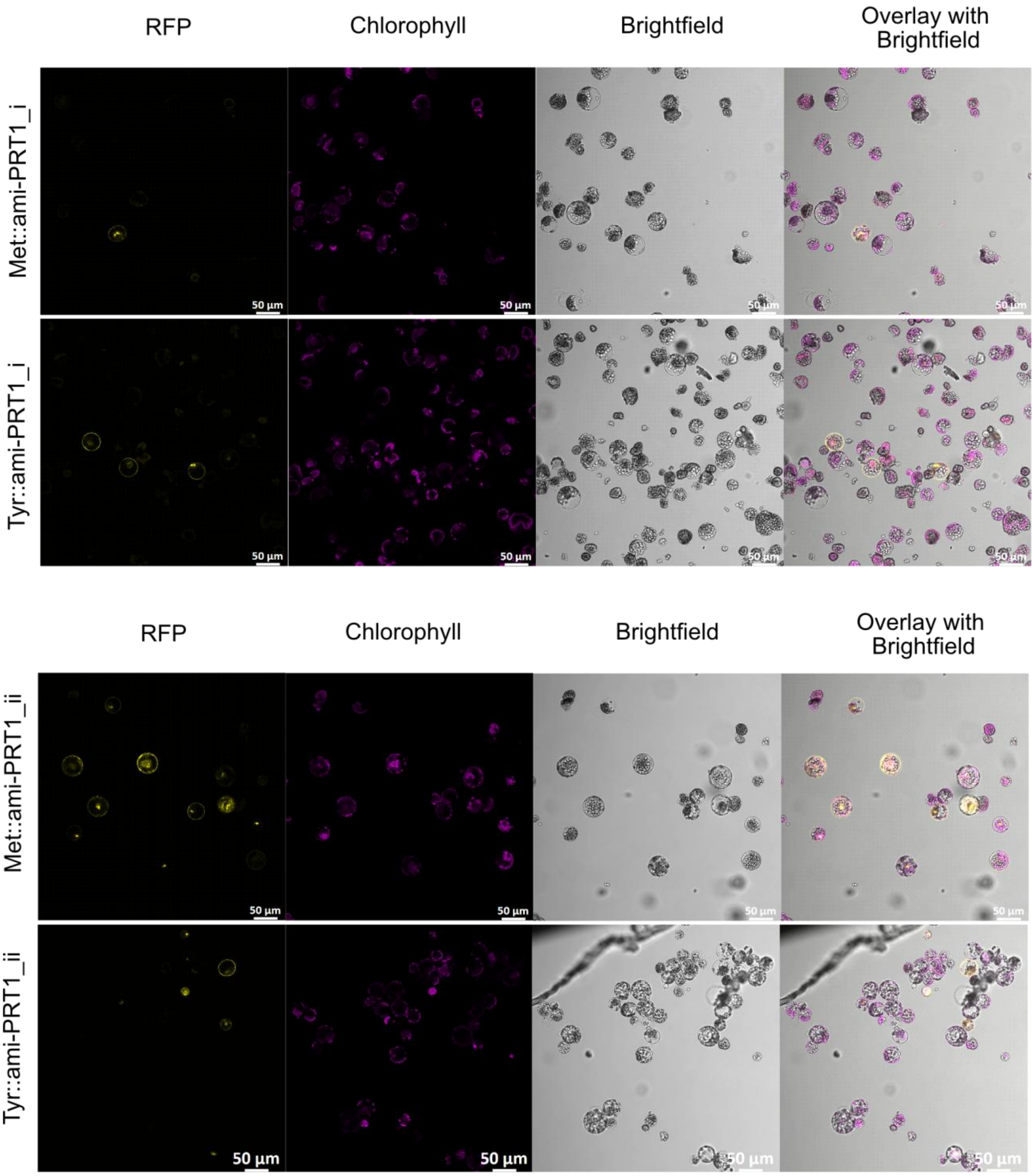


**Supplementary Fig 5.** Representative confocal images of *A. thaliana* protoplasts expressing Met-CBP-RFP and ami-PRT1_i (Met::ami-PRT1_i), Tyr-CBP-RFP and ami-PRT1_i (Tyr::ami-PRT1_i), Met-CBP-RFP and ami-PRT1_ii (Met::ami-PRT1_ii), or Tyr-CBP-RFP and ami-PRT1_ii (Tyr::ami-PRT1_ii). Images were collected at one day post-transfection. RFP fluorescence is false-coloured yellow, chlorophyll autofluorescence is denoted in magenta. Scale bars = 50 µm.

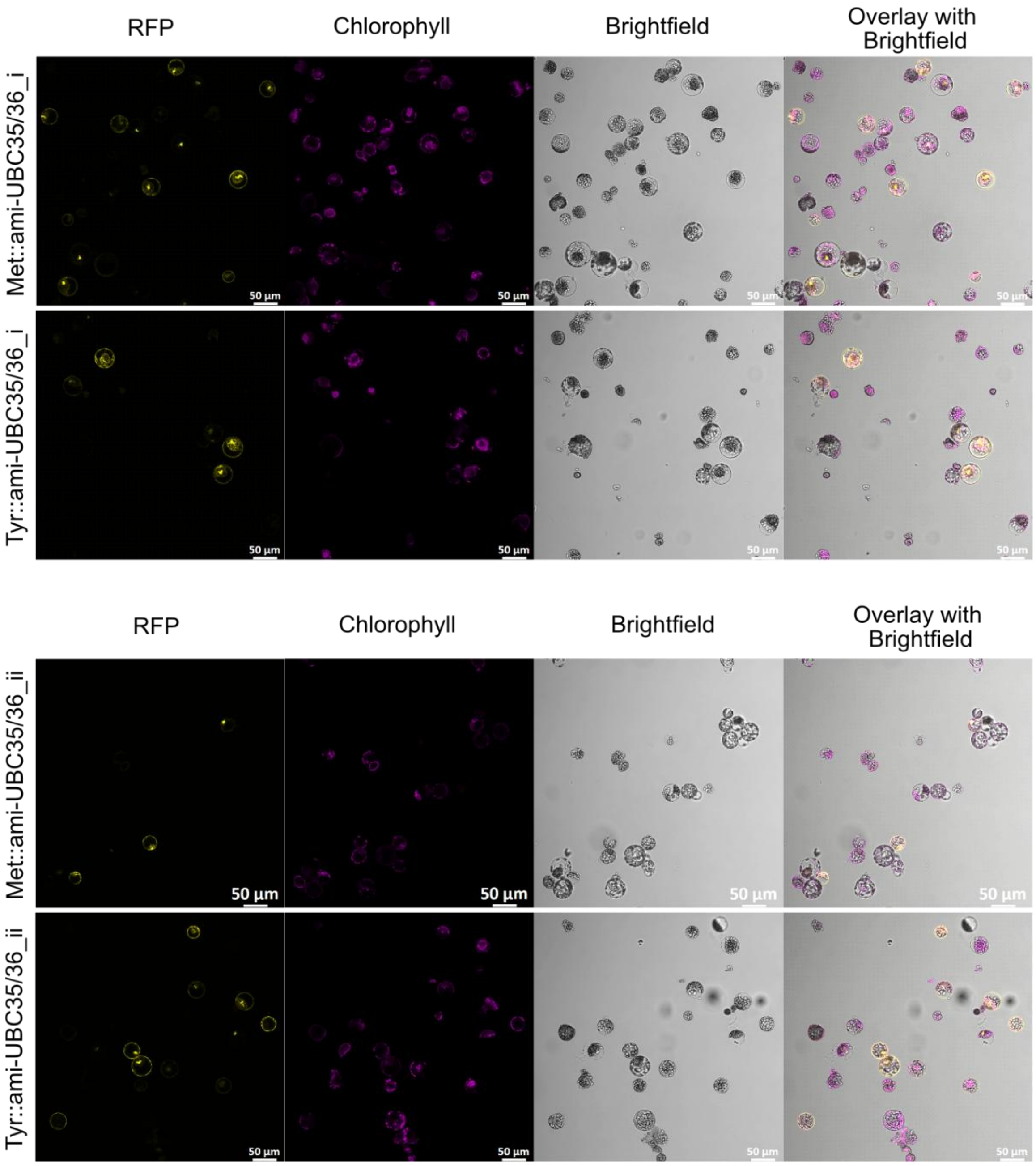


**Supplementary Fig 6.** Representative confocal images of *A. thaliana* protoplasts expressing Met-CBP-RFP and ami-UBC35/36_i (Met::ami-UBC35/36_i), Tyr-CBP-RFP and ami-UBC35/36_i (Tyr::ami-UBC35/36_i), Met-CBP-RFP and ami-UBC35/36_ii (Met::ami-UBC35/36_ii), or Tyr-CBP-RFP and ami-UBC35/36_ii (Tyr::ami-UBC35/36_ii). Images were collected at one day post-transfection. RFP fluorescence is false-coloured yellow, chlorophyll autofluorescence is false-coloured magenta. Scale bars = 50 µm.

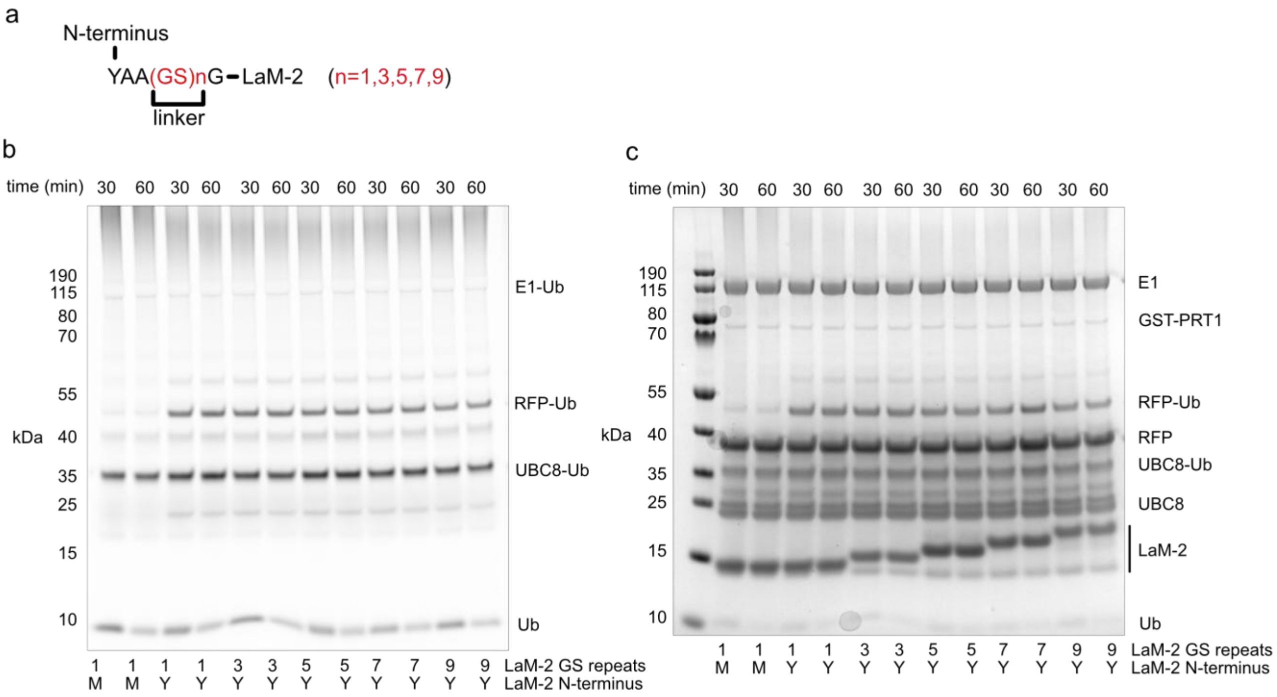


**Supplementary Fig 7. LaM-2 nanobodies with flexible linkers drive ubiquitin transfer to RFP. a** LaM-2 nanobodies with a Tyr N-terminus and between 1 and 9 Gly-Ser (GS) repeats. b GST-PRT1 was incubated with ATP, E1, UBC8, Ub and nanobodies with indicated N-termini (Cy3 fluorescence image). **c**. Coomassie blue stained gel.
